# Scaling-Up Vertical Wheel Bioreactors Based on Cell Aggregate Exposure to Shear Stress and Energy Dissipation Rate

**DOI:** 10.64898/2026.03.24.713990

**Authors:** Julia E. S. Bauer, Faisal J. Alibhai, Pouyan Vatani, David A. Romero, Michael A. Laflamme, Cristina H. Amon

**Affiliations:** Department of Mechanical and Industrial Engineering, University of Toronto, Toronto, Canada; McEwen Stem Cell Institute, University Health Network, MaRS Centre, Toronto, Canada; Department of Laboratory Medicine and Pathobiology, University of Toronto, Toronto, Canada; Institute of Biomedical Engineering, University of Toronto, Toronto, Canada

**Keywords:** Pluripotent Stem Cells (PSCs), vertical wheel bioreactor, *in silico*, shear stress, energy dissipation rate, cell proliferation

## Abstract

**Purpose:** Large quantities of human pluripotent stem cells (hPSCs) are required for clinical applications. 3D suspension cultures are suitable for large scale manufacturing of hPSCs but yield, viability and quality are affected by the hydrodynamic environment. This paper characterizes the hydrodynamic environment inside vertical wheel bioreactors (VWBRs) as a function of size and agitation rates, measures its effect on cell aggregation and proliferation, and proposes the use of Lagrangian-based shear stress and energy dissipation rate (EDR) exposures to support scale-up.

**Methods:** *In silico:* Transient, 3D, turbulent flow simulations are conducted for two VWBR sizes (100, 500 mL) at five agitation rates between 20 and 80 rpm. Trajectories of cell aggregates of sizes from 200 to 1,000 microns are calculated, and shear stress and EDR exposures are collected along these trajectories. *In vitro:* ESI-017 hPSCs were cultured in VWBRs for 6 days. Aggregation efficiency and daily fold ratios were calculated based on cell counts and initial inoculation density.

**Results:** Aggregate size, agitation rate and bioreactor size modulate cell aggregate exposures to EDR and shear stress, which significantly depart from maximum or volume average metrics used for scale-up. Combined *in vitro*/*in silico* results show EDR affects aggregation efficiency, cell counts and aggregate size, and has a small effect on daily fold ratios but a significant effect on total fold ratio.

**Conclusion:** History of trajectory-based cell aggregate exposures to EDRs provide a better scale-up basis for VWBRs than volume-averaged EDR. Shear stress does not significantly affect hPSC aggregation, proliferation and expansion in VWBRs under the tested conditions.

## Introduction

Cardiovascular disease (CVD) is the leading cause of death worldwide[1]. From 1990 to 2019, the number of individuals affected by CVD worldwide increased by 252 million, and it is projected to increase by 3.6% every year from 2025 to 2050[1], [2]. Coronary artery disease (CAD), a type of CVD that involves the narrowing of the coronary arteries, is responsible for one-third to one-half of CVD[3], [4]. CAD can lead to a myocardial infarction resulting in permanent damage to the cardiac tissue[3], [5]. Cardiomyocytes (CMs), the main cell type responsible for force generation and thus cardiac function, are lost during these cardiac events, and this loss of contractile units is a key factor responsible for the decline in heart function following a myocardial infarction[5]. CMs have very limited capacity for regeneration, so injury to cardiac tissue is irreversible[5].

One promising approach to restore the contractile function of the heart is the transplantation of human pluripotent stem cell-derived cardiomyocytes (hPSC-CMs), which stably engraft and electromechanically couple with the host myocardium[6], [7]. Clinical translation of this promising therapy requires scaled manufacturing of hPSC-CMs to generate the very large quantities for even a single dose (∼1 × 10^9^ cells per injured heart)[8]. Traditional manufacturing methods, such as static 2D and 3D culture systems, have had limited success in producing high quantities of human pluripotent stem cells (hPSCs) for differentiation, while minimizing cost and minimally perturbing critical signalling pathways which facilitate CM differentiation[9], [10].

Suspension cultures in bioreactors are increasingly being used to perform stem cell expansion and differentiation processes and are advantageous due to their ability to support larger cell yields while maintaining hPSC pluripotency and enhance oxygen and nutrient circulation[8], [11]. Suspension cultures enable the formation of three-dimensional aggregates and enable recapitulation of mechanical forces which are difficult to obtain in static cultures, such as flow-induced shear stress, which has been shown to modulate signalling pathways through mechano-transduction[12], [13]. One such bioreactor, the vertical wheel bioreactor (VWBR), has recently gained popularity with its U-shaped bottom and impeller design[14]. This design differs from the commonly used stirred tank bioreactor, which has a cylindrical shape and a horizontally rotating impeller[15]. Furthermore, the impeller in the VWBR occupies a larger percentage of space compared to the stirred tank bioreactor, ranging between 22% and 33% depending on VWBR size as opposed to 5% in the stirred tank bioreactor[15]. The VWBR design results in combined radial and axial fluid flow, allowing for more homogeneous mixing and consequently more uniform force distribution compared to the mostly radial mixing in the stirred tank bioreactor[14], [15].

The mechanical environment inside bioreactors can be controlled through changes in operational parameters (e.g., flow rates, agitation rates) and/or changes in geometry. This is particularly important for general purpose dynamic bioreactors, since the optimal mechanical environment is cell-line-dependent, with a wide variety of dose-response curves, albeit these curves are not yet fully quantified[16], [17]. Different bioreactor designs and flow rates are more likely to promote or hinder cell expansion[18]. Human adult mesenchymal stem cells (h-MSCs), for example, have higher final day cell counts when cultured in a perfusion bioreactor under lower flow rate conditions compared to higher flow rate, and h-MSC expansion in hollow fiber bioreactors are associated with low final day cell counts[18]. Mechanical stimuli may initiate cell apoptosis through intrinsic and extrinsic pathways, although the exact dosage and mechanisms are not well understood[19]. In the case of shear stress, the maximum allowable shear stress for various cell types has been reported to vary over two orders of magnitude[17]. For instance, exposure to high shear stress (>0.7 Pa) negatively affects pluripotent stem cell (PSC) viability and proliferation during culture in stirred tank bioreactors[20], [21]. It is important to note that these levels are not typically reached in most research labs expanding PSCs, as the standard operating levels of many bioreactor systems are well below this threshold. However, the shear stress levels experienced during culture within these systems does impact several important aspects of PSC function and cell proliferation, for instance, cell signalling pathways[18], [22]. Cell signalling pathways modulate important cellular processes such as cell maintenance and differentiation, and it is therefore important to understand how these pathways are influenced by the hydrodynamic environment in dynamic culture systems[11]. The most well studied is the impact shear stress has on the signalling pathways responsible for regulating pluripotency and differentiation (e.g. β-catenin and YAP/TAZ signalling)[22], [23], [24]. Shear stress can also influence aggregate size during 3D culture, which can in turn affect access to nutrients within the inner portion of the aggregate. Therefore, tight regulation of shear stress is critical for successful and reproducible expansion and differentiation of PSCs within bioreactor systems.

The growing demand for regenerative therapies urges the need to streamline the transition from concept to clinical applications, a process that can benefit from the efficiency of computational models[16]. Computational Fluid Dynamics (CFD) is a collection of methods for the numerical solution of the system of partial differential equations governing fluid flow[25]. CFD enables high-resolution characterization of fluid-induced forces and mixing inside a bioreactor as a function of its geometry and operational parameters. Importantly, CFD predictions of flow fields, particle transport and mixing dynamics have been validated against power and torque measurements[26], particle image velocimetry (PIV) data[27], [28], and even pulsed laser induced fluorescence (PLIF) [29]on stirred tank bioreactors. Thus, validated CFD models, coupled with *in vitro* cell culture experiments in bioreactors, helps elucidate the impact of fluid-induced forces on the proliferation and differentiation behaviors of hPSCs, and is thus key for the optimization of experimental protocols combining *in silico* and *in vitro* experiments. Furthermore, CFD models are valuable tools to support the design and optimization of bioreactors for scaled-up production of cellular products. By providing inexpensive, *in silico* predictions of flow-induced forces, advection, diffusion and reaction processes, designers can ensure that a consistent mechano-bio-chemical environment can be attained in a variety of bioreactor designs of different sizes.

Clinical applications of novel stem-cell based therapeutics require scaled-up production of cellular products[9]. There is an extensive literature on scale-up strategies based on keeping a certain aspect of the culture environment constant across multiple bioreactor sizes. Commonly used scale-up characteristics have been mixing time, agitation rate, impeller tip speed, shear stress, energy dissipation rate (EDR), nutrient depletion time, power per unit volume, or impeller Reynolds number[17], [30]. Where applicable, characterization of the culture environment inside the bioreactor is made in terms of average or maximum values within the bioreactor. For instance, Dang et al. [31]and Borys et al. [30]studied scale-up of VWBRs and stirred suspension bioreactors using various parameters such as CFD-predicted volume-averaged EDR, and developed empirical scale-up equations for the rotational speed of the impeller across several geometrically similar bioreactor sizes. These works provide guidance on which parameter should be kept constant during bioreactor scale-up to increase the likelihood of achieving cell expansion of high quality and quantity. However, the large uncertainty regarding the mechanisms of cell damage, in addition to the large variation of the hydrodynamic environment inside a bioreactor, limits the success of bioreactor scale-up equations to larger bioreactor volumes[10], [17]. We hypothesize that a detailed understanding of how cultured cells, and the larger-size aggregates they may form, travel throughout the bioreactor and are thus exposed to different doses of mechanical stimuli, can provide a better basis for bioreactor scale-up.

In this paper, we propose the use of the history of cell exposure to mechanical stimuli as a basis for scaling-up VWBRs. In this context we considered cells to have been “exposed” to the shear stress or EDR values at all points that form part of the cell trajectory. Through a combined Eulerian-Lagrangian approach, we fully describe the history of shear stress and EDR experienced by cell aggregates as they move inside the VWBR, for different impeller rotational speeds and cell aggregate sizes. We compare the statistics of shear and EDR exposure to commonly used volume averaged and maximum values. Our results show that the shear stress and EDR experienced by cell aggregates are significantly different from volume average (VA) and maximum shear stress and EDR. Experimental results for hPSC culture in VWBR, in combination with *in silico* results, allow users to identify distributions of EDR and shear stress exposures that lead to larger cell expansion, and clearly indicate a dependence of cell expansion metrics on EDR and, to a lesser extent, shear stress.

## Materials and Methods

### CFD Simulations

The 100 mL and 500 mL VWBR geometric models were created using a combination of SolidWorks and COMSOL Multiphysics 6.2[32]. Dimensions were obtained by direct measurement of the device and product documentation[33]. Physics-controlled meshes were created for both bioreactor sizes using default options, and then further adjusted based on the results of a mesh independence study conducted at the largest rotational speed considered (Figures S2 and S3 in the supplementary material). The selected 100 mL VWBR mesh had a total of 482,758 elements with a minimum element size of 0.288 mm and 5 inflation layers. The 500 mL VWBR mesh had a total of 560,733 elements with a minimum element size of 0.394 mm and 5 inflation layers.

The CFD models for the VWBR were created in COMSOL version 6.2 as two-phase simulations with a free air-liquid interface, which allowed for fluid disturbances on the top of the liquid domain, corresponding to a partially filled VWBR. A single transient, 3D turbulent flow simulation was conducted for each combination of VWBR size, agitation rate, and particle size. Air was used as the fluid above the free surface (gray region in Fig. 1), using the default material properties provided by the software (density of 1.204 kg/m3, dynamic viscosity of 1.814 × 10^-5^ Pa · s). mTeSR1, an incompressible Newtonian fluid, was used as the fluid inside the bioreactor, with a dynamic viscosity of 7.65 × 10^-4^ Pa · s and a density of 993 kg/m3[34]. As shown in Fig. 1, the fluid domain (light blue) was divided into a rotating region containing the impeller wheel, and a stationary domain containing the growth media (liquid) and air (gas), with a defined air-fluid interface. Boundary conditions were imposed as no-slip velocity at all boundaries.

**Fig. 1.**
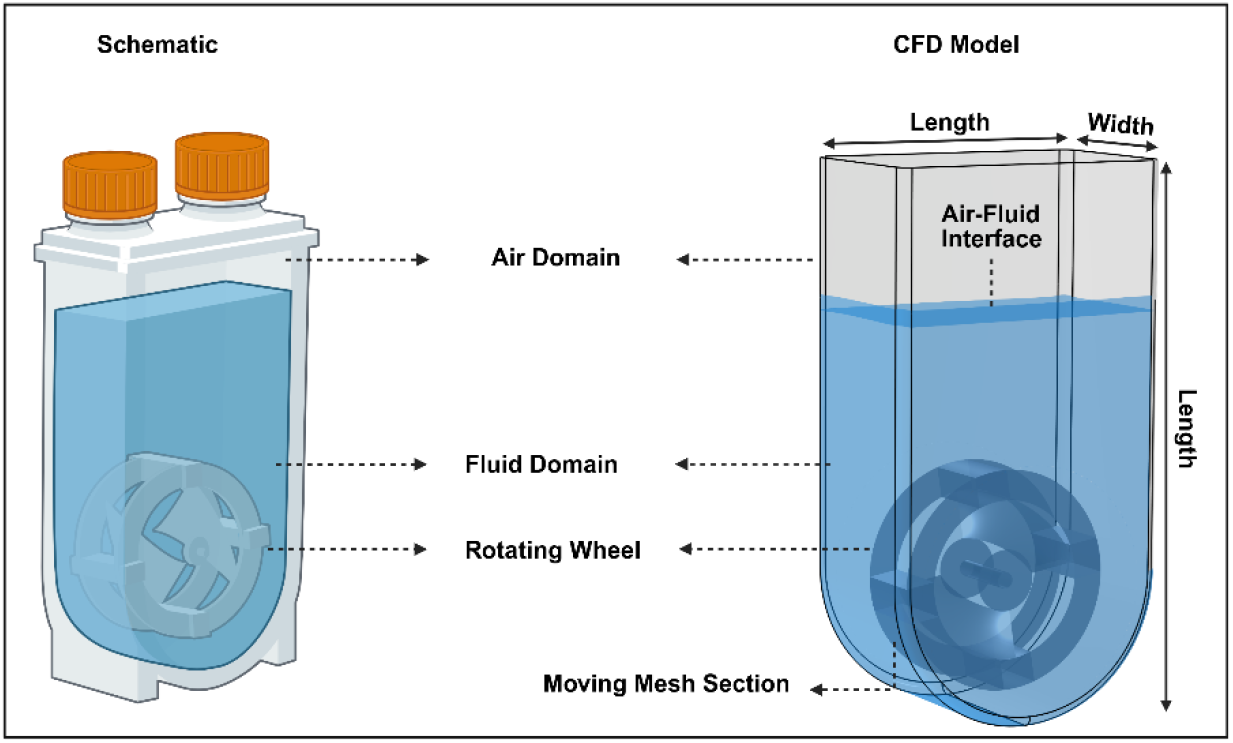
Vertical Wheel Bioreactor (left) and corresponding CFD model (right). Specific dimensions for 100 mL and 500 mL VWBRs are included in Table 1.

Using a constant rotation rate of the impeller wheel, our initial simulation experiments showed that the flow field reaches a steady periodic state after 5 seconds in the 100 mL VWBR and after about 25 seconds in the 500 mL VWBR. Simulation results were stored at a rate of 20 datasets per revolution.

### Governing Equations

The CFD simulations were conducted using the realizable k-ε turbulence model, similar to previous studies[31], [35], with the Reynolds-averaged Navier-Stokes and continuity equations (Equations 1 and 2).

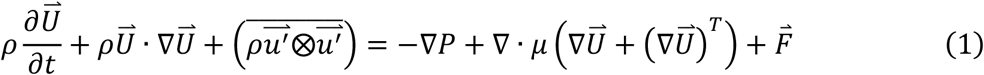

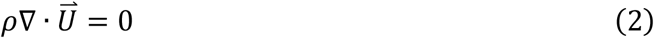

The Reynolds-averaging process separates the flow quantities into average and fluctuating quantities. 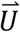 represents the averaged velocity field (m/s) and *u*^′^ represents the fluctuating component (m/s). The Reynolds-averaging process introduces a new term, 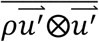, which is known as the Reynolds-stress term.

The Reynolds-stress term is modelled with the Boussinesq hypothesis, outlined in Equation 3.

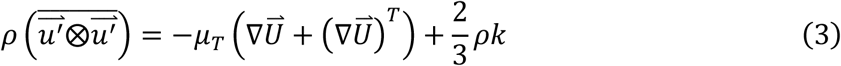

Turbulence closure is introduced with the realizable *k*-ε turbulence model, which solves for the turbulent viscosity, *μ*_*T*_, by introducing two dependent variables, turbulent kinetic energy (*k*) and the turbulent kinetic energy dissipation rate (*ε*), and their respective transport equations. *μ*_*T*_ is described in Equation 4.

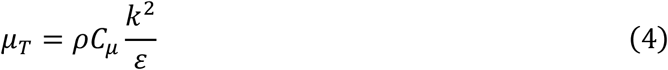

For the realizable *k*-ε turbulence model, *C*_*μ*_ is not a constant and is represented with Equation 5.

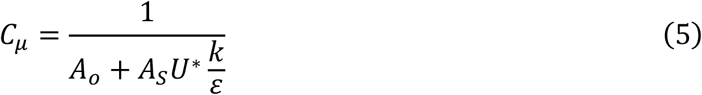

Equations for *A*_*S*_ and *U*^∗^ are presented in Equations 6 and 7 respectively. *A*_*o*_ is a constant and is equal to 4[36].

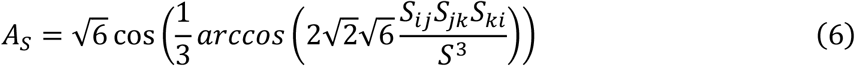

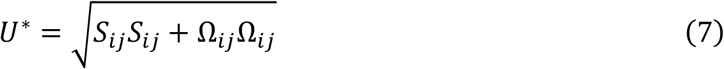

*S, S*_*ij*_, and Ω_*ij*_ are defined in Equations 8, 9, and 10, respectively. *S*_*ij*_ represents the mean strain-rate tensor and Ω_*ij*_ represents the mean rotation-rate tensor.

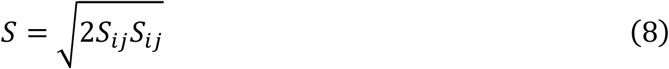

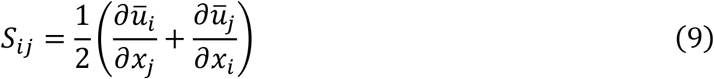

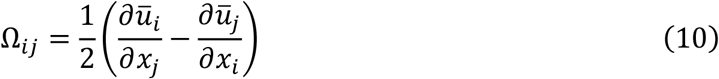

The transport equation for *k* is provided below in Equation 11. *σ*_*k*_ is the turbulent Prandtl number for *k* and has a value of 1 [30]. *P*_*k*_ represents the production term and is given in Equation 12.

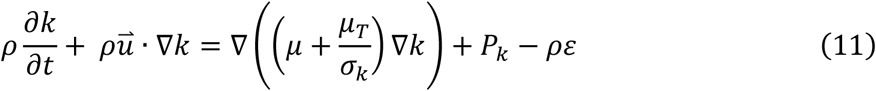

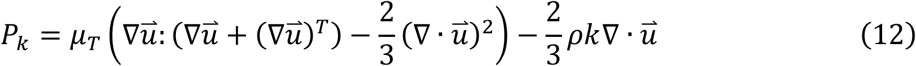

The transport equation for *ε* is given in Equation 13. *σ*_*ε*_ is the turbulent Prandtl number for *ε* and has a value of 1.2[36].

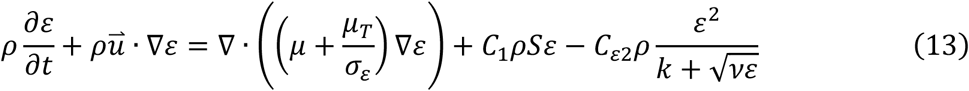

The equation representing *C*_1_ is provided below in Equation 14. *C*_*ε*2_ is a constant with a value of 1.9[36]. *η* is defined using Equation 15.

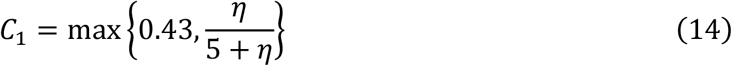

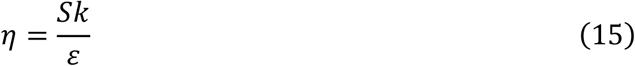

While the realizable k-ε turbulence model effectively captures the mean flow characteristics, as a RANS-based approach, it does not resolve turbulent fluctuations below the Kolmogorov scale. For the specific flow conditions in this study, the Kolmogorov scale falls in the range of 100—250 µm, which is comparable to the size of the aggregates being modelled. The lack of resolution at scales below the aggregate size is considered acceptable for our application, as the main biological effect of the energy dissipation is the modulation of aggregate sizes, and the turbulence power spectra demonstrate that sub-Kolmogorov scales account for only approximately 2% of the total dissipated energy.

Once the mean flow field has been resolved, the energy dissipation rate is available as the solution of Equation 13, and the shear stress can be calculated at any location as the summation of the molecular and turbulent components, i.e.,

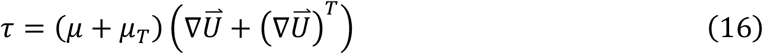

The free surface was modelled using a level set method, described below in Equation 16. *ϕ* represents the level set function. *γ* and *ϵ* are parameters used to ensure numerical stability.

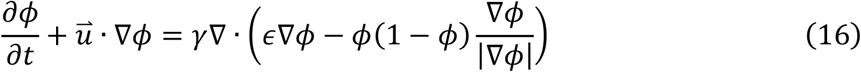

### Cell Aggregate Tracing

Cells are modelled as spherical particles with given mass and diameter. An aggregate, physically consisting of several cells clumped together, is modelled as a single sphere with a diameter that we refer to as the aggregate size. To the authors’ best knowledge, the volumetric density of hPSCs is not available. Thus, te cell aggregate density was assumed to be 1,045 kg/m3, which is representative of the density of adipose tissue mesenchymal stem cells (MSCs)[37]. Both cells and aggregates are referred to in this work as particles, with the terms used indistinctly. In any given simulation, all the aggregates have the same size. This modeling choice was motivated by the need to characterize the shear and EDR exposures of particles of a given size and mass, since these parameters directly influence their trajectories.

Cell aggregate motion under the effect of drag, lift, gravitational, pressure-gradient and virtual mass forces, is modelled using a Lagrangian approach as described below. Basset history forces were neglected in our analysis due to their computational complexity and based on published work that has identified the forces considered here as the most dominant. To calculate particle trajectories in the flow, we integrate Newton’s second law of motion for each particle using a fixed time step, as described in Equation 17. Note that *m*_*p*_ represents the mass of the cell aggregate (kg) and 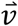 is the cell aggregate velocity (m/s).

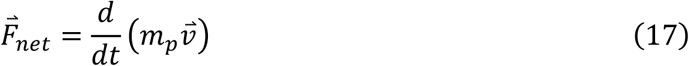

Drag, lift, and gravitational (buoyancy) forces (N) were applied to the cell aggregates, shown in Equations 18, 19, and 20, respectively. The centrifugal force is accounted for with the rotating moving mesh. *d*_*p*_ and *r*_*p*_ are the diameter (m) and radius (m) of the cell aggregates, and *ρ*_*p*_ and *ρ* are the cell aggregate and fluid densities (kg/m3), respectively.

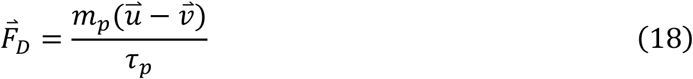

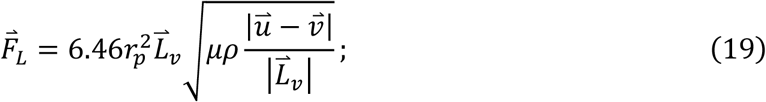

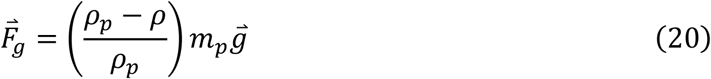

Cell aggregate response time (s), *τ*_*p*_, can calculated using Equation 21, and varies according to relative Reynolds number, *Re*_*r*_, and the drag coefficient, *C*_*D*_. *Re*_*r*_ is described using Equation 22.

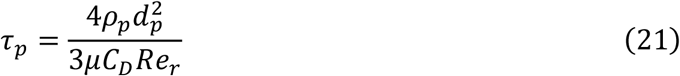

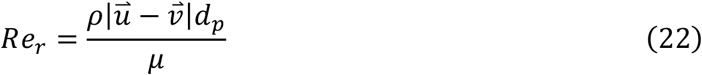

Standard drag correlations were used to compute the drag force, which applies different drag coefficients, *C*_*D*_, depending on the value of the particle relative Reynolds number, *Re*_*r*_, which ranged from *Re*_*r*_ = 10 to *Re*_*r*_ = 25 in our simulations. Since the fluid viscosity *μ* (kg/(m · s)), is low and the cell aggregate size varies, there may be large differences between the fluid velocity (m/s), 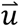, and the particle velocity 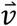, and thus the value of *Re*_*r*_ may vary considerably.

The Saffman lift force was applied to the cell aggregates since the cell aggregates are traveling through a non-uniform velocity field and because the effects of nearby walls can be neglected. Equation 23 describes the vector 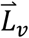. Finally, the gravitational force accounts for the effects of buoyancy with the addition of *ρ* in Equation 20.

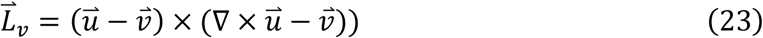

Since the cell aggregate and fluid densities are similar in magnitude, the pressure gradient and virtual mass forces are significant and thus considered in our analysis. The virtual mass force (N) acts on the cell aggregate from the surrounding fluid as the surrounding fluid gets carried with the cell aggregate, virtually increasing the cell aggregate mass[38]. It is presented in Equation 24, where *m*_*f*_ is the mass of fluid displaced by the cell aggregate (kg) and is described in Equation 25. The pressure gradient force (N) is described in Equation 26.

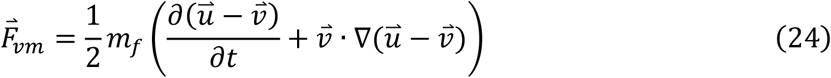

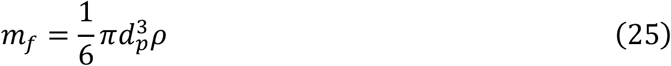

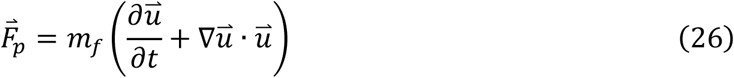

We analyzed a total of 100 equally-sized particles representing cells or cell aggregates with diameters of 200 µm (4.4 µg), 300 µm (15 µg), 400 µm (35 µg), 600 µm (118 µg), 800 µm (280 µg), and 1,000 µm (547 µg). All particles were released at the initial timestep (30 seconds), were randomly placed throughout the mTeSR1 fluid domain and were followed for 30 additional seconds. A statistical study subsampling the collection of 100 particles without replacement, was conducted to assess the independence of the target exposure metrics to the number of particles and the length of time the particles were tracked. Results from this statistical study are included in Figs. S4-S11. Particle-wall interactions were modelled as kinetic-energy-conserving diffuse scattering for all particle sizes. All force calculations were made using the local value of the fluid density, since the presence of a fluid-air interface and the wheel motion induced a vertically varying density profile, as shown in Fig. S15.

### Cell Culture

Undifferentiated ESI-017 (BioTime) hPSCs were thawed from a master cell bank and cultured on human embryonic stem cell (hESC)-qualified Matrigel (Corning) coated tissue culture plastic in mTeSR1 medium (StemCell Technologies). hPSCs were cultured in 2D on Matrigel coated plates for one passage after thaw, after which cells were expanded in 3D using the Vertical Wheel Bioreactor System (PBS Biotech). To inoculate the VWBR on day 0 (D0) of the expansion protocol, 2D cultured hPSCs were dissociated using Accutase and 8 × 10^4^ single cells/mL were seeded into a 100 mL VWBR in mTeSR1 supplemented with 10 µM Y-27632. VWBRs were maintained at either 30, 40, or 60 rpm for the duration of the experiment. Media was exchanged 50% on day 2 and 80% on days 3, 4 and 5 using mTeSR1 by allowing the aggregates to settle and aspirating spent media. No media was exchanged on day 1 (D1). Each day a 1 mL sample was taken from the bioreactor to measure aggregate size and determine cell density within the bioreactor. On day 6, aggregates were harvested from the VWBR and dissociated using Accutase. The final cell density was determined using an automated fluorescent cell counter Cellometer K2 (Revvity). Final day cell counts were divided by the initial inoculation density (D0) or D1 cell counts to determine fold expansions for each condition. Percent aggregation efficiency was defined as the cell count on D1 divided by the initial inoculation density. All cells were cultured in a standard cell culture incubator (5% CO_2_). All hPSC work was approved by the Stem Cell Oversight Committee of the Canadian Institutes of Health Research.

## Results

### Flow Field

Figures 2 and 3 show, respectively, the shear stress and EDR distributions inside the VWBR, and velocity streamlines colored by velocity magnitude. As expected, dark blue regions indicative of low flow velocity, shear stress, and EDR, are mainly located in the volume above the rotating wheel (we refer to this region as the upper domain in the remainder of this paper). Similarly, the highest flow velocity, shear stress and EDR are found near the rotating wheel, specifically surrounding the wheel vanes (Fig. 2). By comparing the flow field between the two bioreactor sizes, it can be observed that shear stress and EDR are larger in the 500 mL VWBR than in the 100 mL VWBR for the same agitation rate. Figure 3 shows that a circulation pattern with two zones develop inside the VWBR, with the primary circulation zone surrounding the rotating wheel, and a secondary circulation zone located in the upper domain. It is also noticeable that that the fluid tends to travel upwards along the walls of the VWBR and downwards along the centre. Although Fig. 3 depicts streamlines for the 100 mL VWBR with a 40 rpm agitation rate, similar flow patterns are observed at all agitation rates in both sizes of the VWBR.

**Fig. 2.**
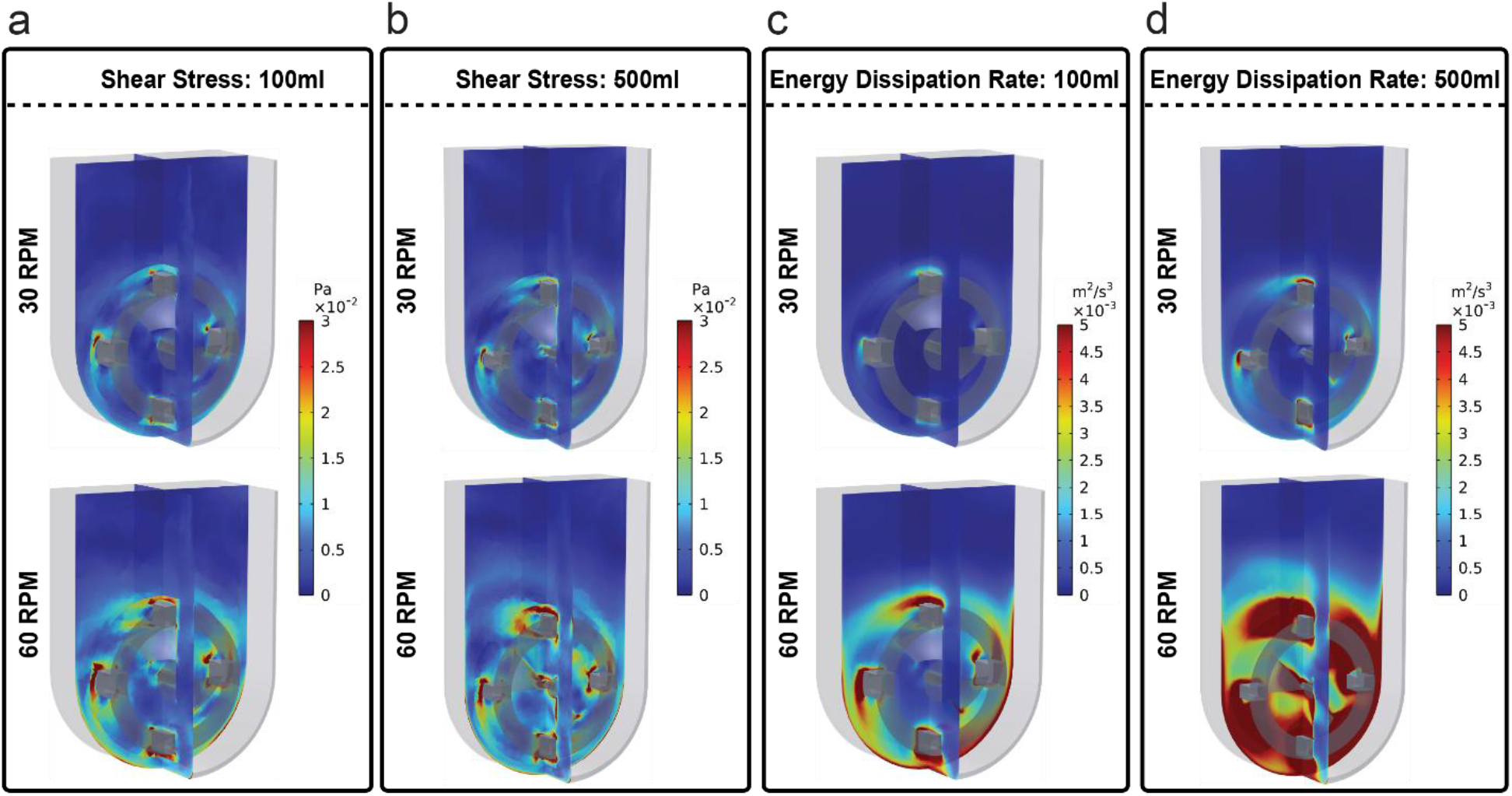
CFD predictions of shear stress and energy dissipation rate (EDR) in VWBR of two sizes at 2 agitation rates. (a) Shear stress (Pa), 100 mL VWBR, (b) shear stress (Pa), 500 mL VWBR, (c) EDR (m^2^/s^3^), 100 mL VWBR, and (d) EDR (m^2^/s^3^), 500 mL VWBR. The panels in the top and bottom row correspond to agitation rates of 30 rpm and 60 rpm, respectively.

**Fig. 3.**
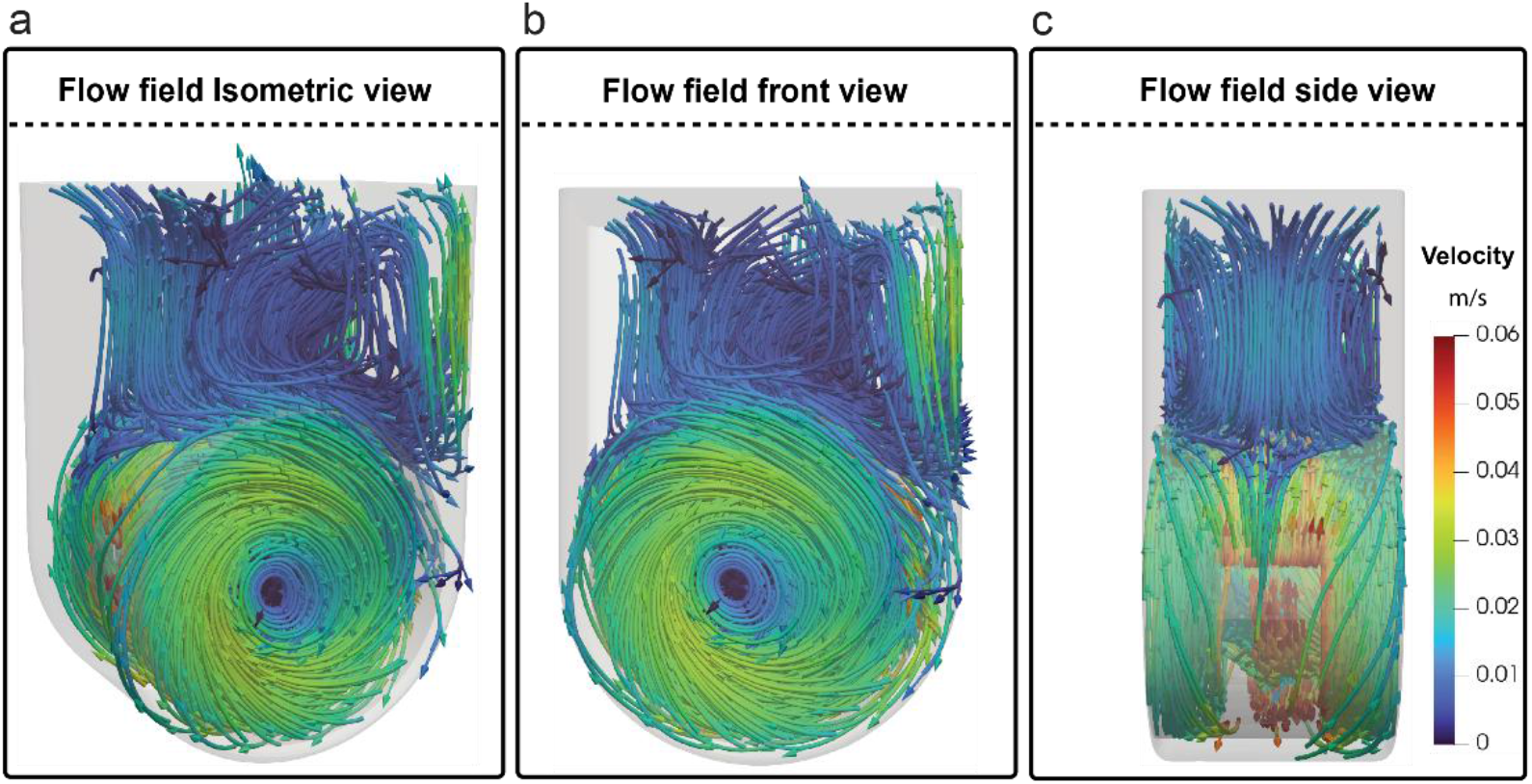
Flow streamlines in the 100 mL VWBR at 40 rpm, colored by velocity magnitude. (a) isometric, (b) front, (c) side views.

### Cell Aggregate Trajectories

Figure 4 shows trajectories for cell aggregates of different sizes, at different (clockwise) agitation rates and in two bioreactor sizes, we will describe the effects of each of these factors separately. As cell aggregate size increases, at a given agitation rate and regardless of bioreactor size, cell aggregate speeds increase and their trajectories change, particularly on the right side of the bioreactor and in the upper domain. For example, in the 100 mL bioreactor at 30 rpm at a cell aggregate size of 547 µg (1,000 µm), many of the cell aggregates in the upper domain are travelling faster than 0.05 m/s, whereas their velocities are less than 0.02 m/s in the upper domain at a cell aggregate size of 4.4 µg (200 µm). Furthermore, larger cell aggregates, such as 547 µg (1,000 µm), move straight downwards, whereas smaller cell aggregates, such as 4.4 µg (200 µm), travel around the upper domain in different directions. Significant differences can also be observed in the maximum height that the cell aggregates reach above the bioreactor wheel. This behaviour is clearly visible across all rows of Fig. 4, where although the cell aggregates may start near the top of the bioreactor, the maximum height of subsequent passes through the upper domain decreases as cell aggregate mass increases.

**Fig. 4.**
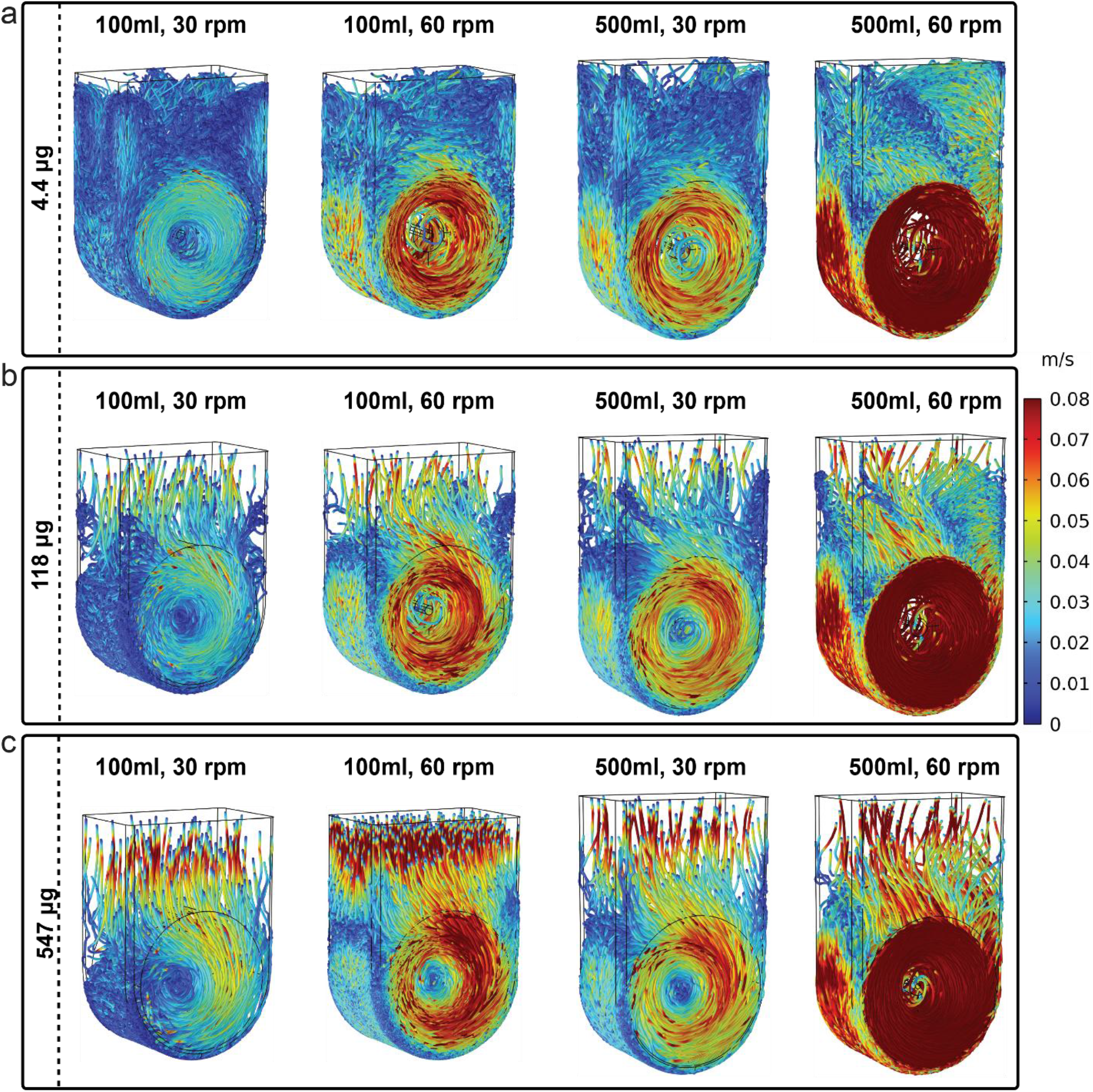
Cell aggregate trajectories, colored by velocity magnitude, for cell aggregates of different sizes at 30 rpm and 60 rpm in the 100 mL and 500 mL bioreactors. Trajectories correspond to 30 seconds of particle motion that initiated after the flow reached steady periodic state.

Regarding the effect of the agitation rate, it can be observed that as agitation rate increases, cell aggregate speeds increase but cell aggregate trajectories remain qualitatively unchanged, except for the larger cell aggregate sizes analyzed in this work. For example, for 118 µg (600 µm) cell aggregates in the 100 mL VWBR (Fig. 4), the cell aggregate paths are essentially the same for 30 and 60 rpm. The main difference between both cases are the cell aggregate speeds, particularly near the wheel. At 30 rpm, the cell aggregate speeds near the wheel obtain a maximum value of approximately 0.05 m/s, whereas at 60 rpm, the maximum cell aggregate speed near the wheel is around 0.08 m/s. In both cases, there are higher cell aggregate speeds in the upper domain. For very large cell aggregates, such as 547 µg (1,000 µm), the cell aggregate paths have less curvature moving downwards at 30 rpm compared to 60 rpm, where the paths still slightly curve to follow the fluid velocity vectors.

Cell aggregates generally travel faster in the 500 mL VWBR as compared to the 100 mL VWBR at the same agitation rate and equivalent cell aggregate size, with exceptions occurring for the larger cell aggregates in the upper domain (Fig. 4). This is expected since at any given agitation rate, a larger wheel radius results in proportionally larger tangential speeds at the outer edges of the wheel. These larger tangential speeds significantly alter cell aggregate trajectories as they transition from the upper domain to the vicinity of the agitation wheel. For example, for 547 µg (1,000 µm) cell aggregates in the 100 mL VWBR, the cell aggregates mainly travel downwards in the upper domain and continue to do so near the wheel whereas the cell aggregate trajectories better align with the fluid velocity vectors in the 500 mL VWBR.

### EDR and Shear Stress on Cell Aggregates

Figure 5 and Fig. 6 show the distributions of EDR and shear stress, respectively, as experienced by 100 cell aggregates (for each cell aggregate size) during their motion inside the VWBR over 30 seconds, corresponding to 10 to 30 wheel revolutions, depending on agitation rate. Note that, for ease of visualization, outliers have been removed. In the figures, the VA EDR and VA shear stress are plotted as blue horizontal lines, while the maximum EDR and maximum shear stress are provided in Table 2. Comparing the boxplots in each subfigure, a clear effect of cell aggregate size becomes apparent. In most cases, cell aggregates of larger sizes experience larger EDR and shear stress, see for example Figs. 5b-c. However, the effect of cell aggregate size is less noticeable as the agitation rate increases, compare for example Fig. 6b with Fig. 6h. A comparison between the left column (100 mL VWBR) with those in the right column (500 mL VWBR) in Fig. 5 and Fig. 6 suggests that at any given cell aggregate size and agitation rate, increasing the bioreactor size generally causes equivalently sized cell aggregates to experience higher EDR and shear stress values. This is, of course, expected since at any given agitation rate, larger bioreactor sizes exhibit larger tangential speeds at the wheel edge.

**Fig. 5.**
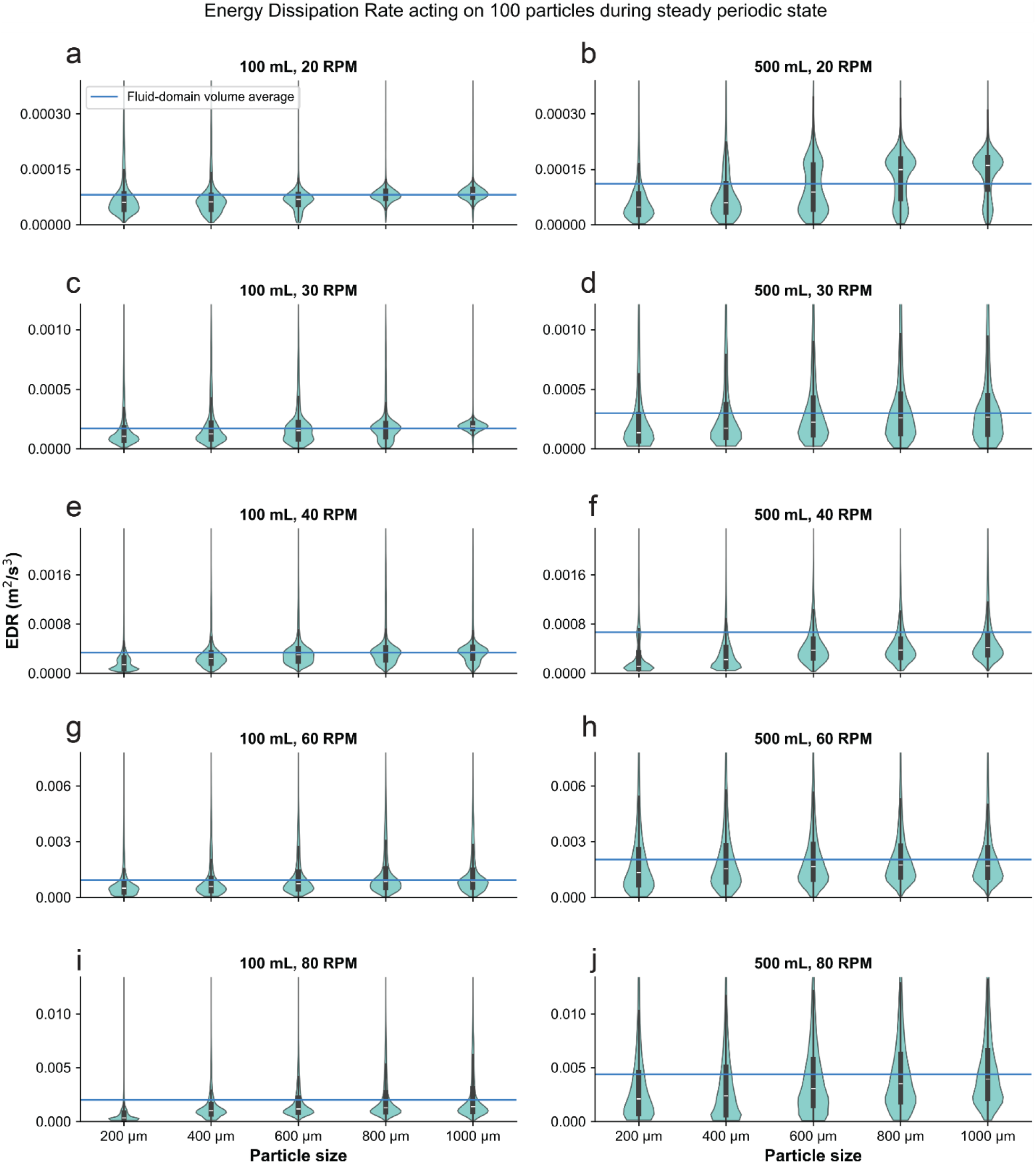
EDR acting on 100 cell aggregates during steady periodic state at various agitation rates and in the 100 mL and 500 mL VWBRs. The blue horizontal line represents the fluid domain volume average (VA) EDR.

**Fig. 6.**
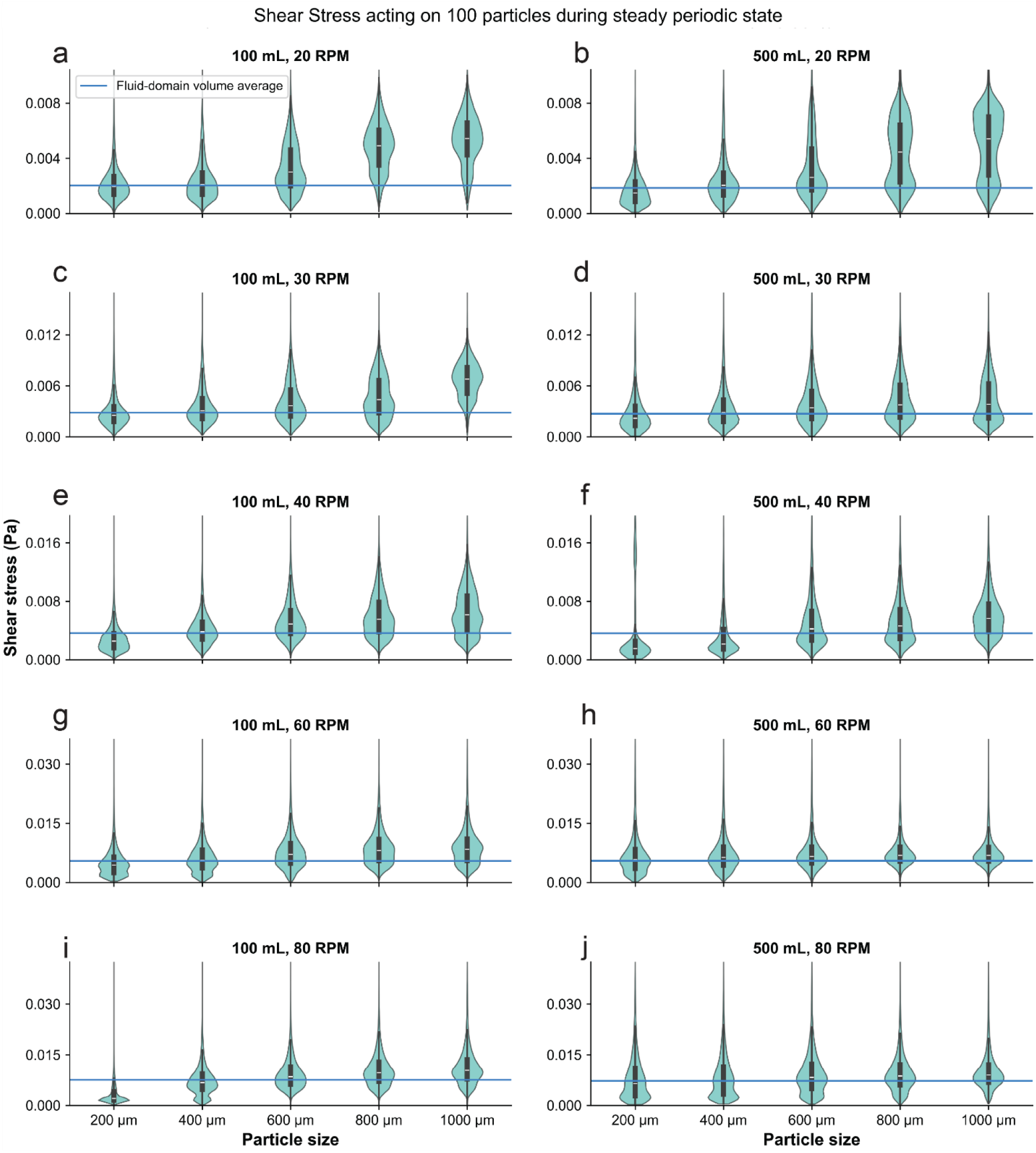
Shear stress acting on 100 cell aggregates during steady periodic state at various agitation rates in the 100 mL and 500 mL VWBRs. The blue horizontal line represents the fluid domain VA shear stress.

Several important observations can be made by comparing the boxplots in Fig. 5 and Fig. 6 with the VA values (horizontal lines in the figures) and maximum values (Table 2). First, none of the tracked cell aggregates encountered the maximum fluid domain shear stress or EDR values (Table 1) over the course of the simulation time, suggesting that characterizing the mechanical environment inside the VWBR in terms of maximum shear stress or maximum EDR may significantly overestimate the actual cell exposure to this environment. Second, all boxplots depict distributions that are skewed towards lower values (right-skewed) but with large tails, indicating that a significant portion of cell aggregate-times are spent at shear stress and EDR values that are significantly above the VA. Specifically, in 42 out of 100 distributions depicted in Fig. 5 and Fig. 6 the median of the distribution is above the VA. Overall, the computed EDR and shear stress distributions indicate that volume-averaged and order statistics (e.g. maximum) are poor predictors of cell aggregate exposure to these mechanical stimuli.

**Table 1.** Overall dimensions of 100 mL and 500 mL vertical wheel bioreactors considered in this study.

|  | 100 mL |  | 500 mL |  |
| --- | --- | --- | --- | --- |
|  | mTeSR | Air | mTeSR | Air |
| Length (mm) | 52.5 | 52.5 | 90.0 | 90.0 |
| Width (mm) | 32.5 | 32.5 | 50.0 | 50.0 |
| Height (mm) | 75.0 | 25.0 | 145.0 | 30.0 |
| Filled volume (mL) | 68.6 |  | 355.4 |  |

**Table 2.**
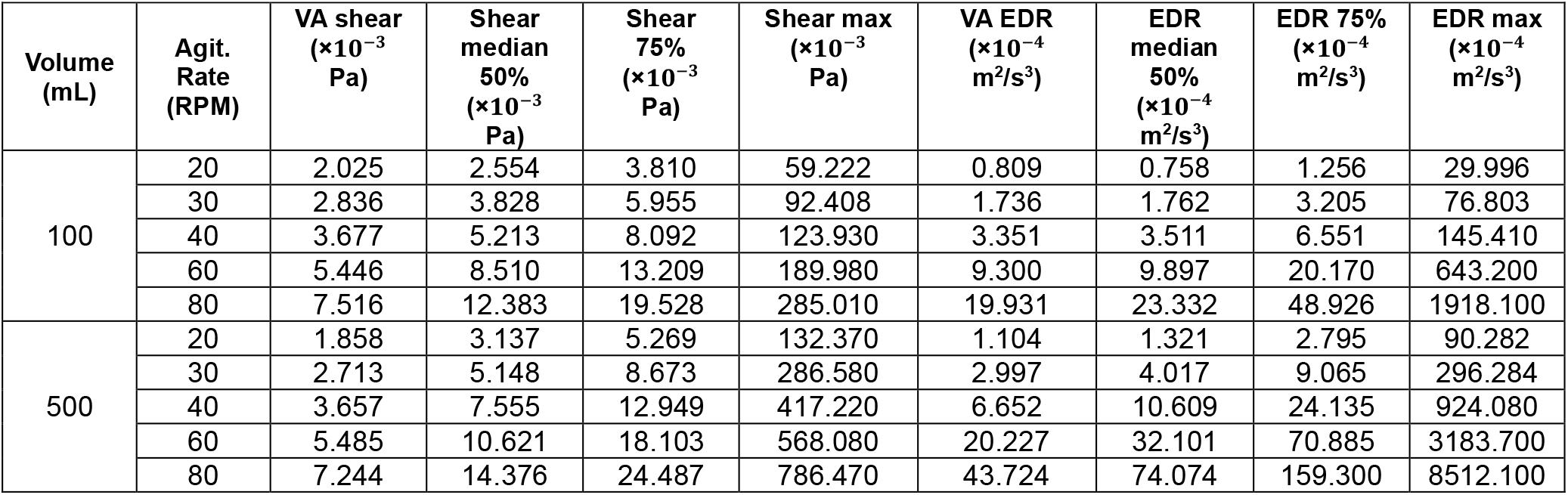
Eulerian-based shear stress and EDR order statistics for different agitation rates and VWBR sizes.

| Volume (mL) | Agit. Rate (RPM) | VA shear ($\times 10^{-3}$ Pa) | Shear median 50% ($\times 10^{-3}$ Pa) | Shear 75% ($\times 10^{-3}$ Pa) | Shear max ($\times 10^{-3}$ Pa) | VA EDR ($\times 10^{-4}$ m<sup>2</sup>/s<sup>3</sup>) | EDR median 50% ($\times 10^{-4}$ m<sup>2</sup>/s<sup>3</sup>) | EDR 75% ($\times 10^{-4}$ m<sup>2</sup>/s<sup>3</sup>) | EDR max ($\times 10^{-4}$ m<sup>2</sup>/s<sup>3</sup>) |
| --- | --- | --- | --- | --- | --- | --- | --- | --- | --- |
| 100 | 20 | 2.025 | 2.554 | 3.810 | 59.222 | 0.809 | 0.758 | 1.256 | 29.996 |
|  | 30 | 2.836 | 3.828 | 5.955 | 92.408 | 1.736 | 1.762 | 3.205 | 76.803 |
|  | 40 | 3.677 | 5.213 | 8.092 | 123.930 | 3.351 | 3.511 | 6.551 | 145.410 |
|  | 60 | 5.446 | 8.510 | 13.209 | 189.980 | 9.300 | 9.897 | 20.170 | 643.200 |
|  | 80 | 7.516 | 12.383 | 19.528 | 285.010 | 19.931 | 23.332 | 48.926 | 1918.100 |
| 500 | 20 | 1.858 | 3.137 | 5.269 | 132.370 | 1.104 | 1.321 | 2.795 | 90.282 |
|  | 30 | 2.713 | 5.148 | 8.673 | 286.580 | 2.997 | 4.017 | 9.065 | 296.284 |
|  | 40 | 3.657 | 7.555 | 12.949 | 417.220 | 6.652 | 10.609 | 24.135 | 924.080 |
|  | 60 | 5.485 | 10.621 | 18.103 | 568.080 | 20.227 | 32.101 | 70.885 | 3183.700 |
|  | 80 | 7.244 | 14.376 | 24.487 | 786.470 | 43.724 | 74.074 | 159.300 | 8512.100 |

**Table 3.** Concordance analysis correlating meaningful changes in biological outcomes with statistically significant differences in hydrodynamic indicators. Indications of True or False refer to whether the metrics in the corresponding column are significantly (p < 0.05) or meaningfully different between the compared conditions. Eulerian-based VA EDR are always different, so this column contains the % differences in their values.

| Comparisons | Lagrangian EDR Exposures | Eulerian VA EDR | Aggregation Efficiency | Aggregate Size | Final/D0 Fold Ratio | Final/D1 Fold Ratio | Final/D2 Fold Ratio |
| --- | --- | --- | --- | --- | --- | --- | --- |
| 100 mL 30 rpm vs. 100 mL 40 rpm | False | 93% | False | True | False | True | True |
| 100 mL 30 rpm vs. 100 mL 60 rpm | True | 435.7% | True | True | False | True | True |
| 100 mL 30 rpm vs. 500 mL 30 rpm | True | 72.6% | True | True | False | True | True |
| 100 mL 40 rpm vs. 100 mL 60 rpm | True | 177.5% | True | True | False | True | True |
| 100 mL 40 rpm vs. 500 mL 30 rpm | False | 11.8% | True | True | False | True | True |
| 100 mL 60 rpm vs. 500 mL 30 rpm | True | 210.3% | True | True | False | True | True |

### Experimental Results

Boxplots of the final day cell counts and aggregate sizes at different agitation rates in the 100 mL and 500 mL bioreactors are presented in Fig. 7. It can be observed that the agitation rate modulates final day aggregate sizes, with 40 rpm resulting in the smallest variance of aggregate sizes. Importantly, a 40-rpm agitation rate also results in larger final day cell counts than 30 rpm or 60 rpm. These results suggest that 40 rpm would be a suitable agitation rate for hPSC culture in a 100 mL VWBR. Scaling based on a Eulerian metric, such as volume-averaged (VA) shear stress, would suggest using an agitation rate of 30 rpm in the 500 mL bioreactor to achieve approximately the same VA shear, 5.2 × 10^−3^ Pa. However, experimental tests of this scaling scenario shown in Fig. 7 indicate that it yields significantly lower final cell counts, a similar median aggregate size, but a very different aggregate size distribution. As we will see in the remainder of this section, *in silico* results provide a richer description as a basis for optimizing the cell culture protocol.

**Fig. 7.**
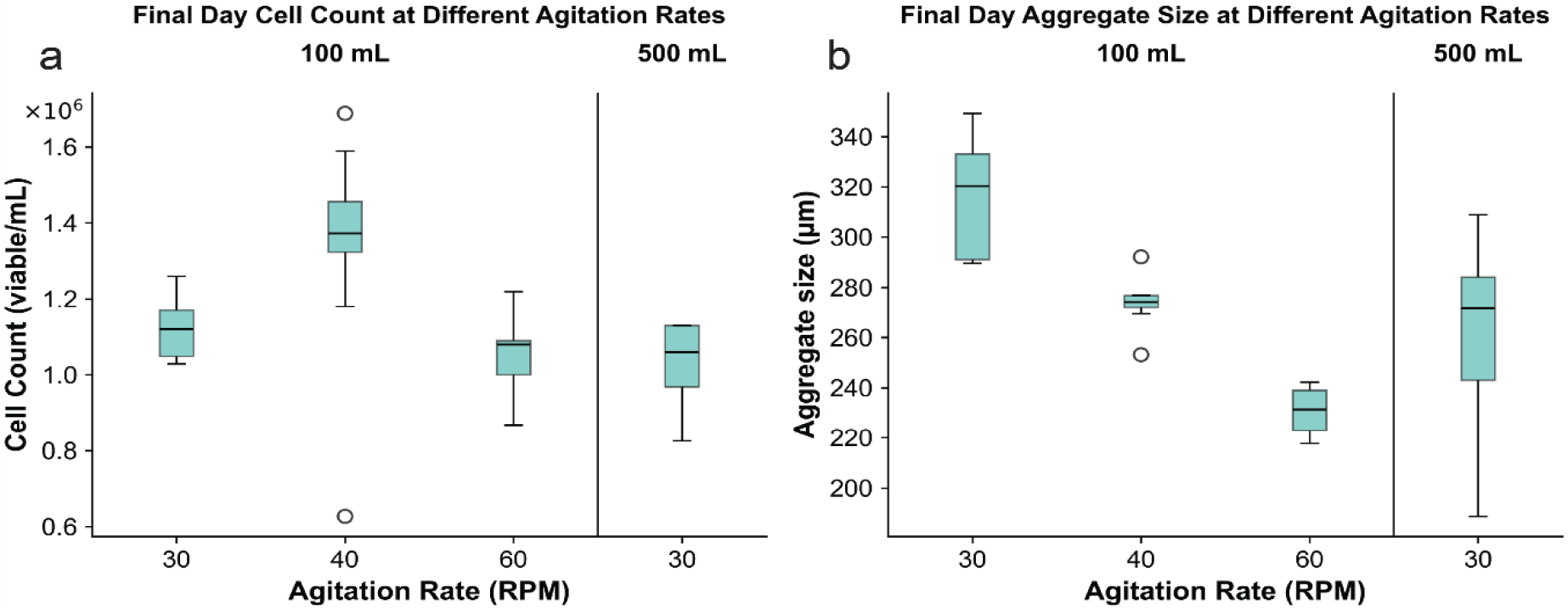
Cell proliferation and aggregation results at different agitation rates in the 100 mL and 500 mL VWBR. (a) Final day cell counts and (b) final day aggregate sizes.

Figure 8 shows bivariate boxplots that consider experimental outcomes, such as final day cell counts, final day aggregate sizes, cell count fold ratios, and aggregation efficiency, as a function of the distribution of shear stress and EDR experienced by the cell aggregates. We note that a few outliers have been removed from this figure for ease of visualization. EDR and shear stress results from cell aggregates with a diameter of 300 µm were selected as this represented the average final day aggregate size in our experimental data. VA values are depicted in the figures as vertical dashed lines with the same color as the experimental data corresponding to the same agitation rate. Note that the VA EDRs for all three agitation rates are closer to the 75^th^ percentile of the EDRs experienced by the aggregates, whereas the VA shear stresses are closer to the median of the Lagrangian-based shear stress distribution.

**Fig. 8.**
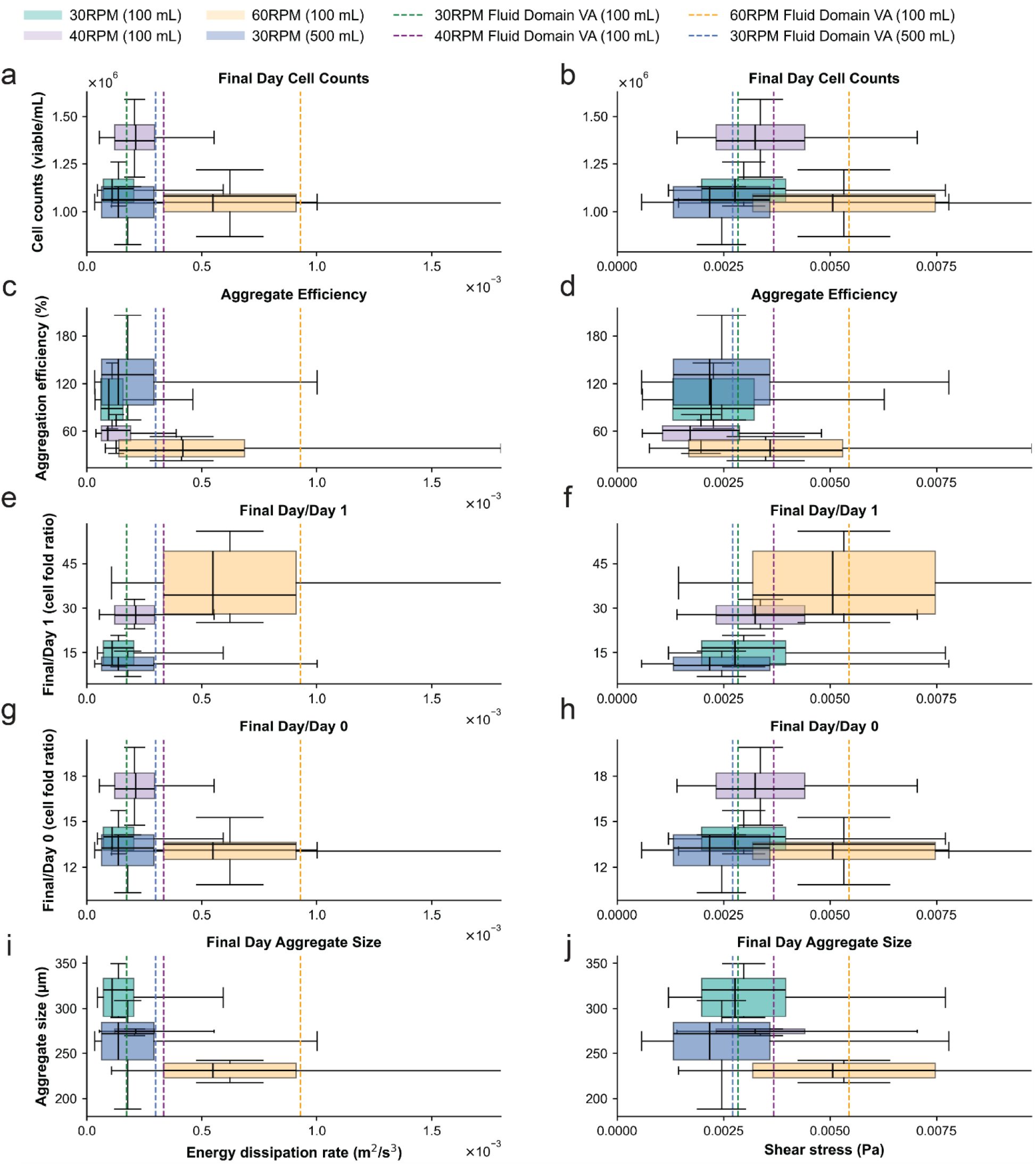
(a,b) Final day cell counts, (c,d) aggregate efficiency, (e,f) final day/D1 fold ratio, (g,h) total (final day/D0) fold ratio, and (i,j) aggregate size as a function of EDR (left column) and shear stress (right column) distributions at various agitation rates in the 100 mL and 500 mL VWBR. Panels (c) and (d) correspond to D1 cell and cell aggregate sizes of 0.55 µg (100 µm) for 30 rpm, 0.28 µg (80 µm) for 40 rpm, and 0.07 µg (50 µm) for 60 rpm. All other panels correspond to final day aggregate sizes of 15 µg (300 µm).

Figures 8a-b compare the final day cell counts for 3 different agitation rates. It can be observed that the highest final day cell counts are obtained at EDR < 3.0 × 10^−4^ m^2^/s^3^and *τ* < 5.0 × 10^−3^ Pa, corresponding to an agitation rate of 40 rpm. However, the differences in final day cell counts cannot be explained solely by EDR or shear stress differences, as their distributions overlap horizontally for the 30 rpm and 40 rpm cases. In addition, the final day cell counts for 40 rpm and 60 rpm vertically overlap despite a significant difference in EDR and shear distributions. Aggregation efficiency, i.e., the percentage of seeded cells that aggregate and survive the first day of culture, is shown in Figs. 8c-d. In this case, the highest aggregation efficiency is observed for EDR < 2.0 × 10^−4^ m^2^/s^3^ and *τ* < 3.0 × 10^−3^ Pa, corresponding to an agitation rate of 30 rpm. Note the aggregation efficiencies for 40 rpm and 60 rpm are significantly lower, reaching a median of 40% at 60 rpm. Figures 8e-f and g-h show respectively, final day/D1 and final day/D0 cell count fold ratios. It is observed that EDR and, to a lesser extent, shear stress have a significant impact on fold ratios, with an agitation rate of 60 rpm resulting in the largest fold ratios after D1, corresponding to 2.5 × 10^−4^ m^2^/s^3^ < EDR < 9.0 × 10^−4^ m^2^/ s^3^, and 3.0 × 10^−3^ Pa < *τ* < 8.0 × 10^−3^ Pa. Note, however, that the lower fold ratio observed for 40 rpm is compensated with a larger aggregation efficiency on D0, thus the largest cell counts on the final day of culture are obtained at 40 rpm.

Finally, aggregate size distributions are shown in Figs. 8i-j. A clear effect of EDR and shear stress are observed, with little vertical overlap between the boxes. Note that a 40-rpm agitation rate results in the lowest variance of aggregate sizes, and that both 30 rpm and 40 rpm result in aggregate sizes in the range of 280-340 µm, which in our experience are optimal for PSCs at this stage. As expected from Fig. 7 results, Figs. 8i-j show overlapping aggregate size distributions for 500 mL at 30 rpm and 100 mL at 40 rpm.

Overall, Fig. 8 clearly indicates that EDR and shear stress have a significant effect on the formation of cell aggregates in the earlier stages of the process, while they have a smaller but cumulative effect on cell proliferation, at least for the ranges of EDR and shear stress studied in this work. Results also show that EDR modulates all outcomes of interest to a greater extent than shear stress does. Importantly, the finding that agitation rates have different effects on aggregation and proliferation suggests that variable-speed protocols may be necessary for an optimal culture.

## Discussion

Bioreactor scale-up equations are important to ensure that similar, if not equivalent, mechanical environments can be obtained in geometrically similar bioreactors of different sizes. Traditional scale-up methods use either 1) an Eulerian-based approach, where volume-averages, order statistics or integrals of the flow field, such as power density, torque, velocity magnitude, shear stress, or EDR are kept constant between bioreactor scales, or 2) a kinematic approach, where the impeller tip speed or agitation rate is used to drive scale-up decisions[17], [30]. The major limitation of these approaches is their lack of consideration to the forces experienced by the cell aggregates during bioreactor operation. Forces such as drag, lift, centrifugal and gravitational forces act on the cell aggregates as they travel throughout the fluid domain and cannot be accounted for when analyzing the fluid domain alone. These forces, which are dependent on the cell aggregate size and density, impact the cell aggregate trajectories throughout the bioreactor, and result in the cell aggregates experiencing shear stress and EDR that may differ significantly from VA and maximum values of shear stress and EDR.

Inside the VWBR, the relative magnitude of drag, lift, centrifugal and gravitational forces acting on the cell aggregates, and their interaction with the flow field, dictate the speeds and trajectories of the cell aggregates and, consequently, their exposure to shear stress and EDR. Bioreactor size, agitation rate and cell aggregate mass significantly affect these forces and thus cell aggregate speeds and trajectories. Cell aggregate speed generally increases if two of these parameters are kept constant and the other parameter increases. Firstly, when agitation rate and bioreactor size are held constant but cell aggregate mass increases, the gravitational force becomes the dominant force at larger cell aggregate sizes and therefore increases the downwards speeds of the cell aggregates. Secondly, when the cell aggregate mass and bioreactor size are held constant, an increase in agitation rate causes an increase in the fluid inertial force throughout the entire bioreactor, resulting in higher cell aggregate speeds. Finally, when agitation rate and cell aggregate size are kept constant but bioreactor size increases, cell aggregates generally travel faster due to the increased fluid inertial force corresponding to larger tangential velocities with the larger 500 mL bioreactor wheel.

Cell aggregate trajectories, on the other hand, change in response to variations in cell aggregate and bioreactor size but are less influenced by agitation rate. When agitation rate and bioreactor size are held constant but cell aggregate mass increases, cell aggregate trajectories increasingly deviate from the velocity vectors due to larger inertial and gravitational forces. As agitation rate increases, the fluid inertial force and centrifugal forces also increase, both acting to dominate over the gravitational force, resulting in cell aggregate trajectories that closely align with the velocity vectors. Larger bioreactor sizes result in unchanged trajectories near the wheel with significant changes in trajectories in the upper domain, where inertial, gravitational and centrifugal forces are more balanced.

Due to the differences in cell aggregate trajectories with respect to changes in agitation rate, bioreactor size, and cell aggregate mass, the resulting history of cell aggregate exposures to EDR and shear stress are affected by these parameters as well (Fig. 5 and Fig. 6). Increases in agitation rate increase the EDR and shear stress experienced by the cell aggregates due to larger velocity and velocity gradients near the bioreactor wheel. A similar effect is observed when comparing two bioreactor sizes at most agitation rates and cell aggregate masses, again due to increased tangential velocity at the bioreactor wheel due to its larger size in the 500 mL VWBR. As cell aggregate size increases, gravitational forces start to dominate and cause the cell aggregates to spend more time around the bioreactor wheel where EDR and shear stress are highest. However, this trend is partially damped at larger agitation rates due to a changing balance between inertial and gravitational forces and the presence of two circulation zones (Figs. 3 and 4). Finally, IQR of the EDR and shear stress distributions are typically lower for smaller cell aggregate sizes (4.4 µg, 35 µg, and 118 µg) than for larger cell aggregates (280 µg, 547 µg), and this trend is observed for both bioreactor sizes and at almost all agitation rates. This suggests decreased variability of EDR and shear stress exposure, and this is consistent with smaller cell aggregates travelling more in the secondary circulation zone found in the upper domain.

The cell aggregate trajectories analyzed in this work clearly show that, during the observation period (10 to 30 wheel revolutions), cell aggregates do not travel throughout all the fluid domain and, consequently, a Lagrangian-based analysis provides a better representation of the mechanical environment experienced by the aggregates than Eulerian-based volume-averaged metrics. For instance, Figs 9a-b are cases where the VA EDR is within the operating range suggested by Dang et al.[31], represented by red and blue lines in the figure. Note that data represented in each of these boxplots is calculated based on cell aggregate trajectories with a fixed time step, and thus the portions of the distributions outside these recommended ranges map directly to times of exposure to these potentially damaging EDRs. Consider, for example, 4.4 µg (200 µm) cell aggregates in each case presented in Fig. 9. Even though both conditions (100 mL at 60 rpm and 500 mL at 30 rpm) have the same VA EDR, the EDRs experienced by equivalently sized cell aggregates are different. Namely, for 100 mL at 60 rpm (Fig. 9a), the 4.4 µg (200 µm) cell aggregates encounter EDRs that are outside the recommended VA EDR range approximately 25% of the time (7.5 s), while this increases to 75% of the time (22.5 s) in the 500 mL VWBR at 30 rpm. To the best of our knowledge, the mechano-transduction dynamics of hPSCs has not been thoroughly characterized. Hence, we cannot comment on whether these exposure times would trigger signalling cascades that would modulate differentiation fates in subsequent targeted cultures, as has been observed in MSCs that upregulated osteogenic gene expression when exposed to intermittent shear stress ranges similar to those calculated in this work (1.0 × 10^−3^ Pa to 1.0 × 10^−2^ Pa)[39].

**Fig. 9.**
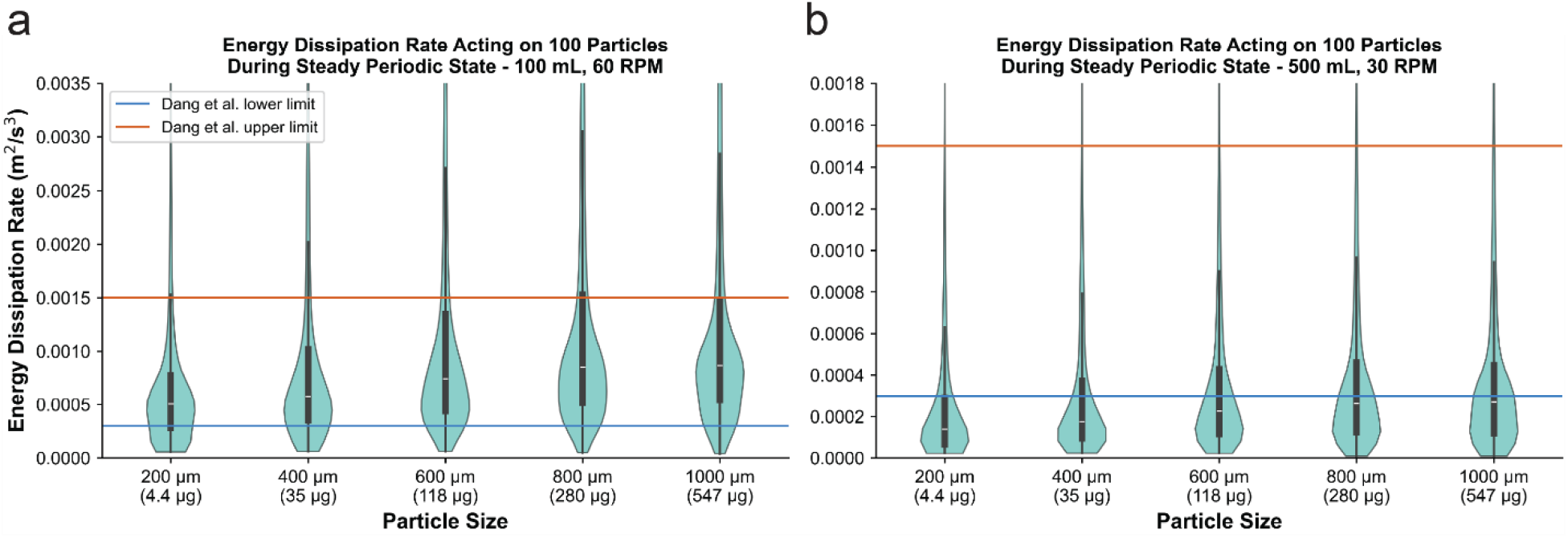
EDR acting on cell aggregates of various sizes compared to the suggested operating range VA EDR values proposed by Dang et al. [31]for (a) 100 mL at 60 rpm and (b) 500 mL at 30 rpm. The red and blue lines represents the upper and lower limits, respectively, of the suggested operating range VA EDR values outlined by Dang et al. [31].

Overall, the preceding discussion leads to the conclusion that volume-averaged or order statistics such as maximum or median values over the fluid domain are poor predictors of cell exposure to mechanical stimuli in a VWBR. This was also confirmed by comparing in silico and in vitro results for two conditions (100 mL at 40 rpm, 500 mL at 30 rpm) that would be considered equivalent according to the VA shear stress. Complex flow features, cell aggregate trajectories, and the resulting exposure to EDR and shear stress cannot be accurately estimated without considering cell size and density. Analysis of EDR and shear stress distributions experienced by cell aggregates of different sizes would be even more important when scaling-up production of PSCs to bioreactors that are geometrically dissimilar.

Experimental results for hPSC suspension culture in VWBR augmented with *in silico* shear stress and EDR cell exposure were presented in Fig. 8 as a function of shear stress and EDR, quantities that describe the mechanical environment that the fluid flow exerts on the cells and are modulated by the agitation rate. The observed variations in aggregate sizes, final day cell counts, fold ratios and aggregation efficiencies suggest the cell expansion process is sensitive to changes in agitation rate, a finding confirmed by previous work[30], [31]. In this work, we explain this effect through changes in the distributions of shear stress and EDR experienced by the cells as they are carried by the flow throughout the bioreactor at different agitation rates. It is observed that shear stress and EDR distributions are very sensitive to agitation rate, cell aggregate size, and bioreactor size, and vary over a wide range, typically spanning one order of magnitude (2X – 10X), for any combination of these factors.

To more formally evaluate which hydrodynamic characterization approach—Eulerian volume-averaged values or Lagrangian particle-tracked distributions—better predicts biological outcomes, we performed a concordance analysis across all valid pairwise condition comparisons (6 comparisons from 4 conditions: 100 mL and 500 mL bioreactors at the available RPM conditions; Table 2). For each condition pair, we assessed whether Lagrangian EDR distributions differed significantly (Mann-Whitney U test, p < 0.05 with false discovery rate corrections), calculated Eulerian volume-averaged EDR differences (range: 11.8% to 435.7%), and examined whether biological outcome metrics exceeded empirically-determined thresholds for meaningful change. These thresholds were set as 30% difference in aggregation efficiency, 20 µm difference in aggregate sizes, and 1.0 difference in fold ratios. A striking pattern emerged: Final/D2 (long-term cell proliferation) differed in all six condition pairs (100% concordance), regardless of whether Lagrangian distributions were significantly different or not. This complete concordance suggests that the Final/D2 fold ratio is determined by factors beyond the current hydrodynamic characterization, or at least that subtle aspects of the hydrodynamic environment not captured by distribution comparisons are critical. In contrast, Eulerian volume-averaged EDR values varied substantially across comparisons (11.8% to 435.7%), yet showed no systematic relationship to Final/D2 changes, i.e., large Eulerian differences coincided with Final/D2 changes in some cases but not others. Aggregation efficiency showed more nuanced behavior: changes were observed in 5 of 6 comparisons, including cases with both significant and non-significant Lagrangian distributions. In 1 of 6 comparisons, no meaningful changes were observed in aggregation efficiency despite a 93% difference in VA EDR. Overall, this suggests that aggregation is influenced by multiple factors.

By considering flow-induced forces on cell aggregates of different sizes in a Lagrangian context, our experimental results augmented with simulation results clearly show that EDR and shear stress distributions, within the ranges studied herein, have a significant effect on the formation of aggregates in the early stages of culture, but have a lesser but still significant impact on cell proliferation and daily fold ratios. This aligns with previous work by Borys et al., 2020, who noted the aggregate size distributions varied across agitation rates whereas there was no significant difference between growth kinetics at different agitation rates[40]. However, our results lead to a different conclusion. Figure 8 shows that agitation rates do have a mild effect on growth kinetics that, though seemingly negligible when analyzing daily fold ratios (Figs. S12 and S13), becomes more significant as a cumulative effect over several days (Figs. 8e-f). Furthermore, both Borys et al., 2018 and Dang et al., 2021 discussed that EDR is a better scale-up basis than shear stress due to its larger influence on aggregate size[30], [31]. This is consistent with our observation of the larger effect of EDR compared to shear stress on aggregation efficiency, cell proliferation and aggregate size.

One limitation of the results presented in this work is that the simulations considered cell aggregates of a single size as they travel for a period of time within a flow field in a steady periodic state. However, cell aggregates progressively grow during the culture period. A more detailed picture of the history of cell exposure to mechanical stimuli could potentially be estimated by combining results for different cell aggregate sizes. For example, shear stress and EDR distributions of single-cell or small multicell sized cell aggregates (such as 50 µm) can be used to represent exposure during initial inoculation and aggregation (D0-D1), results for smaller cell aggregates (100 µm or 200 µm) can be used to estimate cell exposures from days 1 to 3, and medium-sized cell aggregates (300 µm or 400 µm) can be used to estimate shear stress and EDR exposure during days 4 to 6. This can guide dynamic culture protocols, in which the agitation rate is changed over the course of the cell culture period to promote relatively constant shear stress and EDR exposure while minimizing aggregate pooling at the bottom of the bioreactor. This is especially important for high density culture systems due to the increased risk of aggregate fusion[41]. Based on our results, a suitable protocol for the 100 mL bioreactor would be to have a 30-rpm agitation rate for the first day of culture (D0-D1) and then increase the agitation rate to 40 rpm-60 rpm for the rest of the culture period (days 1 to 6), possibly with a ramp schedule.

The dependence of shear stress and EDR distributions on cell aggregate mass and density indicates that not all cell aggregate sizes may be suitable for use in the VWBR. For example, brain organoids derived from hPSCs have been grown under constant agitation in bioreactors and reached 3 mm in diameter, which extrapolated from our simulation results, would likely remain near the bottom of the bioreactor and consequently experience higher shear stress and EDR[42]. It may be that, once these organoids reach a certain size, it would be better to continue their culture process in a different vessel to ensure they do not become damaged from shear stress or EDR.

Another limitation of our results is that cell aggregate collisions were not considered and, thus, aggregate formation and growth was not modelled. Incorporating the physical parameters that characterize bond formation and energy transfer between colliding cells would allow us to associate the mechanical environment, predicted *in silico*, with experimental results on aggregate formation and breakage. Note, however, that such undertaking would require more sophisticated turbulence closures, or perhaps directly resolving (rather than modelling) turbulence features at scales much smaller than the target aggregate sizes. Finally, although mechano-transduction of shear stress and EDR stimuli is known to modulate gene expression and cell signalling pathways, these were not characterized in our *in silico* or *in vitro* work[22], [23]. Consideration of cell signalling pathways would provide a richer biological understanding of the response of PSCs to the suspension culture environment.

In this work, we compared Eulerian-based vs. Lagrangian-based hydrodynamic indices for scaling-up vertical wheel bioreactors (VWBRs). Cell aggregate trajectories in 100 mL and 500 mL VWBRs were described for cell aggregates of various sizes and at different agitation rates. Local shear stress and energy dissipation rates (EDR) experienced by the cell aggregates were collected along their trajectories and represented as distributions. Analysis shows that cell aggregate speeds and trajectories are influenced by drag, gravitational, lift, and centrifugal forces, and change with respect to agitation rates, cell aggregate size, and bioreactor size. Features of the shear stress and EDR distributions, such as median and interquartile range, were observed to be modulated, each to a different extent, by changes in agitation rate, cell aggregate size and bioreactor size. Overall, results show that volume-averaged metrics (e.g., volume average (VA) EDR, VA shear stress) or maxima are not sufficient to describe the mechanical environment the cells are exposed to, particularly when considering different cell aggregate sizes. Experimental results show a stronger influence of EDR on final day cell aggregate size, final day cell counts, and aggregation efficiency, as compared to shear stress. Notably, examination of daily fold ratios showed significant cumulative effects of EDR in contrast to previous work that found growth kinetics to not be significantly affected by it. Overall, our results support the use of Lagrangian-based analyses of cell aggregate trajectories and associated exposures for scale-up studies. This is particularly important in multi-day suspension cultures, where increasing aggregate sizes significantly affect their exposure to shear stress and EDR stimuli, and when conducting scale-up studies in geometrically dissimilar bioreactors.

## Acknowledgements

We acknowledge the support of the Government of Canada’s New Frontiers in Research Fund (NFRF) [NFRFT-2022-00447], the Canada Research Chairs Program (CRC-2020-00245), and the Natural Sciences and Engineering Research Council of Canada (NSERC).

## Statements and Declarations

### Ethics Approval

All human pluripotent stem cell work was approved by the Stem Cell Oversight Committee of the Canadian Institutes of Health Research.

