## Supplementary Material for "Scaling-Up Vertical Wheel Bioreactors Based on Cell Aggregate Exposure to Shear Stress and Energy Dissipation Rate"

Journal Name: Annals of Biomedical Engineering

### Time step independence

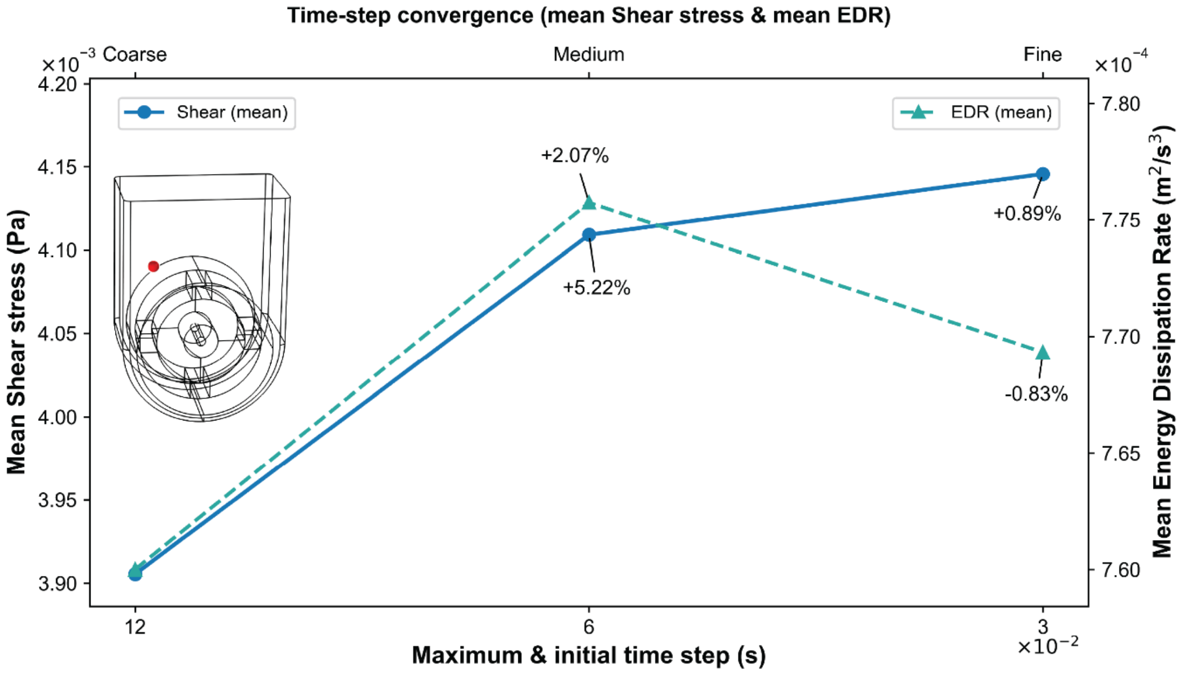


***Fig. S1*** *Mean shear stress (left y-axis) and mean EDR (right y-axis) are reported for coarse, medium, and fine maximum/initial time-step sizes (bottom x-axis; values shown as ×10⁻² s). Means were computed over the last two rotations of the wheel in each simulation. Percentage annotations show the stepwise percent change with respect to the previous (larger) time step, demonstrating convergence of both metrics with time-step refinement*

### Mesh independence analysis (100 mL VWBR)

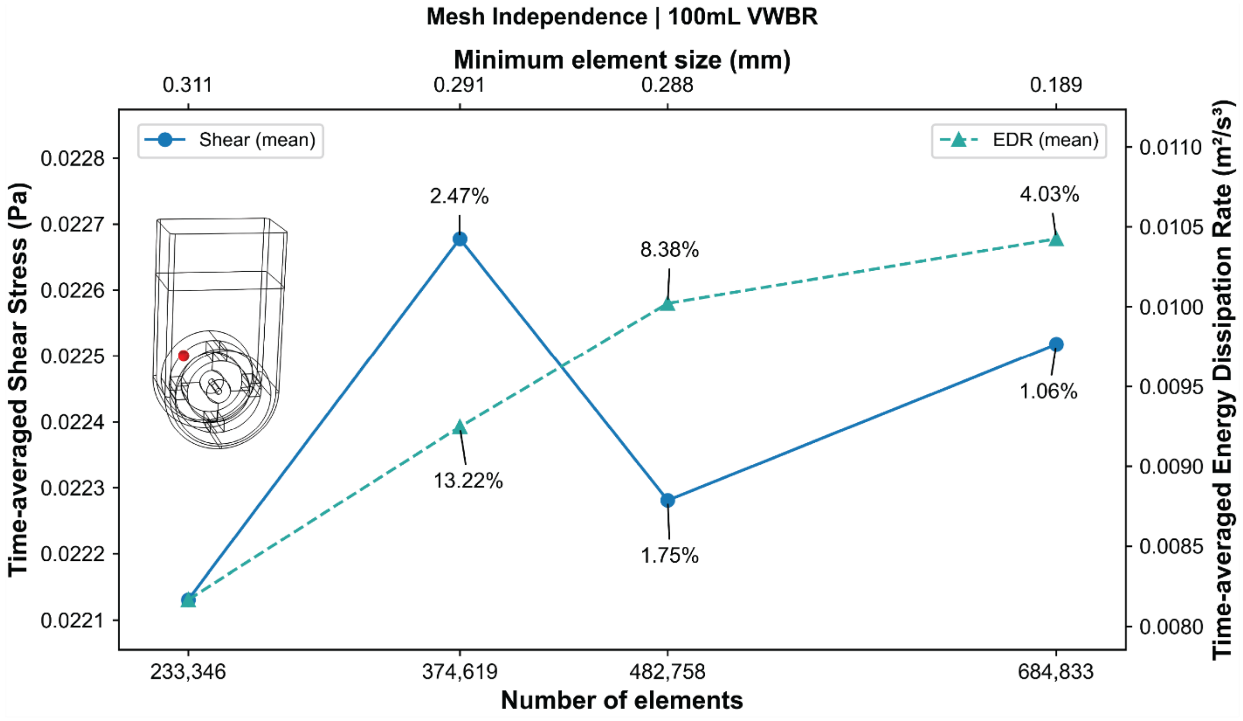


***Fig. S2*** *Mesh independence assessment for the 100 mL vertical wheel bioreactor (VWBR). The mean time-averaged shear stress (left y-axis) and mean time-averaged energy dissipation rate, EDR (right y-axis; $m^2/s^3$), were computed over the last three wheel rotations at the selected probe location for progressively refined meshes (bottom x-axis: number of elements). The corresponding minimum element size for each mesh is shown on the top x-axis. Percentage annotations indicate the stepwise percent change in each metric relative to the previous (coarser) mesh.*

#### Mesh independence analysis (500 mL VWBR)

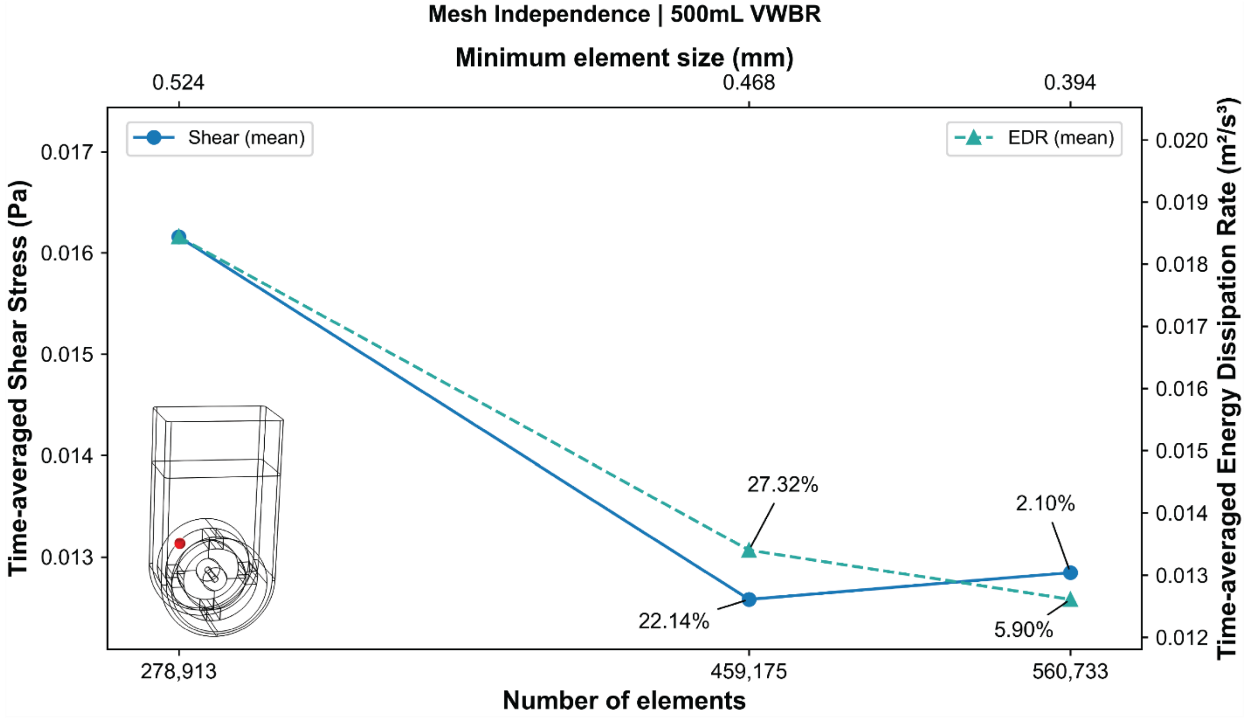


***Fig. S3*** *Mesh independence assessment for the 500 mL vertical wheel bioreactor (VWBR). The mean time-averaged shear stress (left y-axis) and mean time-averaged energy dissipation rate, EDR (right y-axis; $m^2/s^3$), were computed over the last three wheel rotations at the selected probe location for progressively refined meshes (bottom x-axis: number of elements). The corresponding minimum element size for each mesh is shown on the top x-axis. Percentage annotations indicate the stepwise percent change in each metric relative to the previous (coarser) mesh.*

#### Particle number independence (EDR, 100 mL VWBR, 20 rpm)

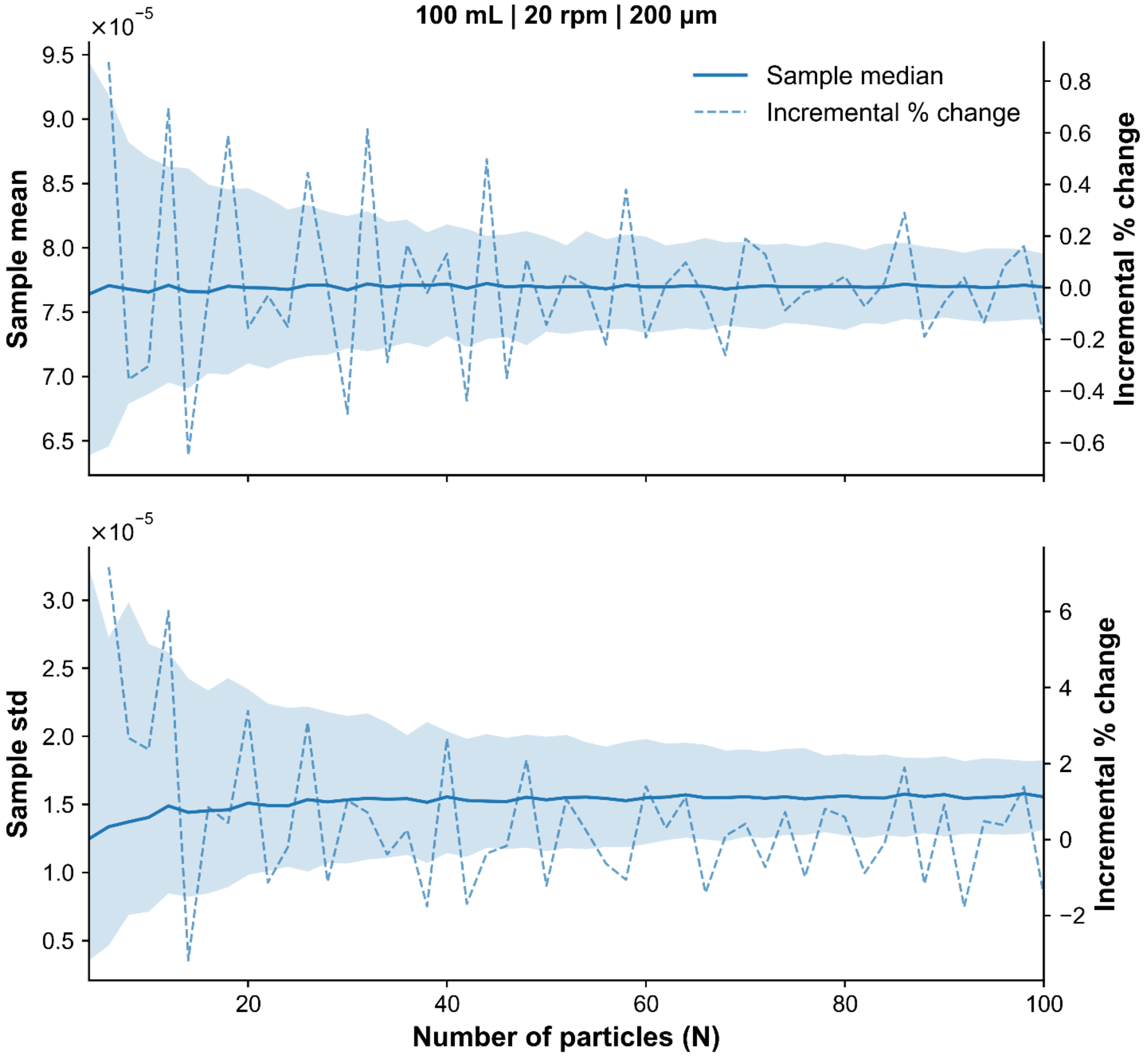


***Fig. S4*** *Particle-number (N) convergence of time-averaged EDR exposure statistics for the 100 mL vertical wheel bioreactor simulated at 20 rpm with 200 μm particles. The solid blue line (median) and shaded band (95% interval) show the across-particle mean (top) and standard deviation (bottom). The 95% interval narrows rapidly with increasing N and is tight by N ≈ 30–50 (very tight at N = 100), supporting 100 particles as a conservative sample size. The dashed curve (right axis) shows the incremental % change between successive N values* $[100\,(y_j - y_{j-1})/y_{j-1}]$*, which fluctuates around zero and diminishes as N increases.*

#### Particle number independence (Shear, 100 mL VWBR, 20 rpm)

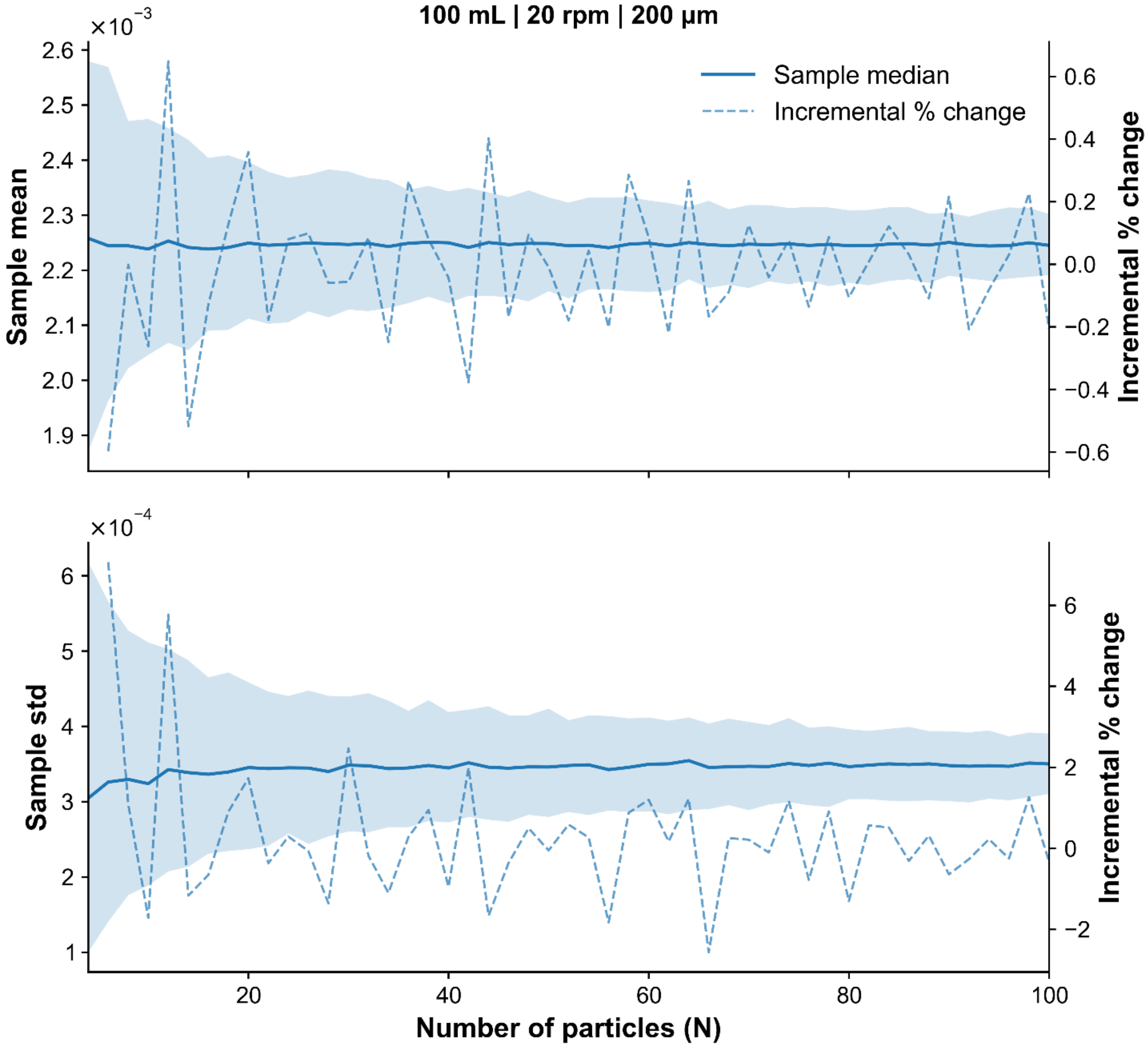


***Fig. S5*** *Particle-number (N) convergence of time-averaged shear-stress exposure statistics for the 100 mL vertical wheel bioreactor simulated at 20 rpm with 200 μm particles. The solid blue line (median) and shaded band (95% interval) show the across-particle mean (top) and standard deviation (bottom). The 95% interval narrows with increasing N and is tight by N ≈ 30–50 (very tight at N = 100), supporting 100 particles as a conservative sample size. The dashed curve (right axis) shows the incremental % change between successive N values* $[100\,(y_j - y_{j-1})/y_{j-1}]$*, which fluctuates around zero and diminishes as N increases.*

#### Particle number independence (EDR, 500 mL VWBR, 80 rpm)

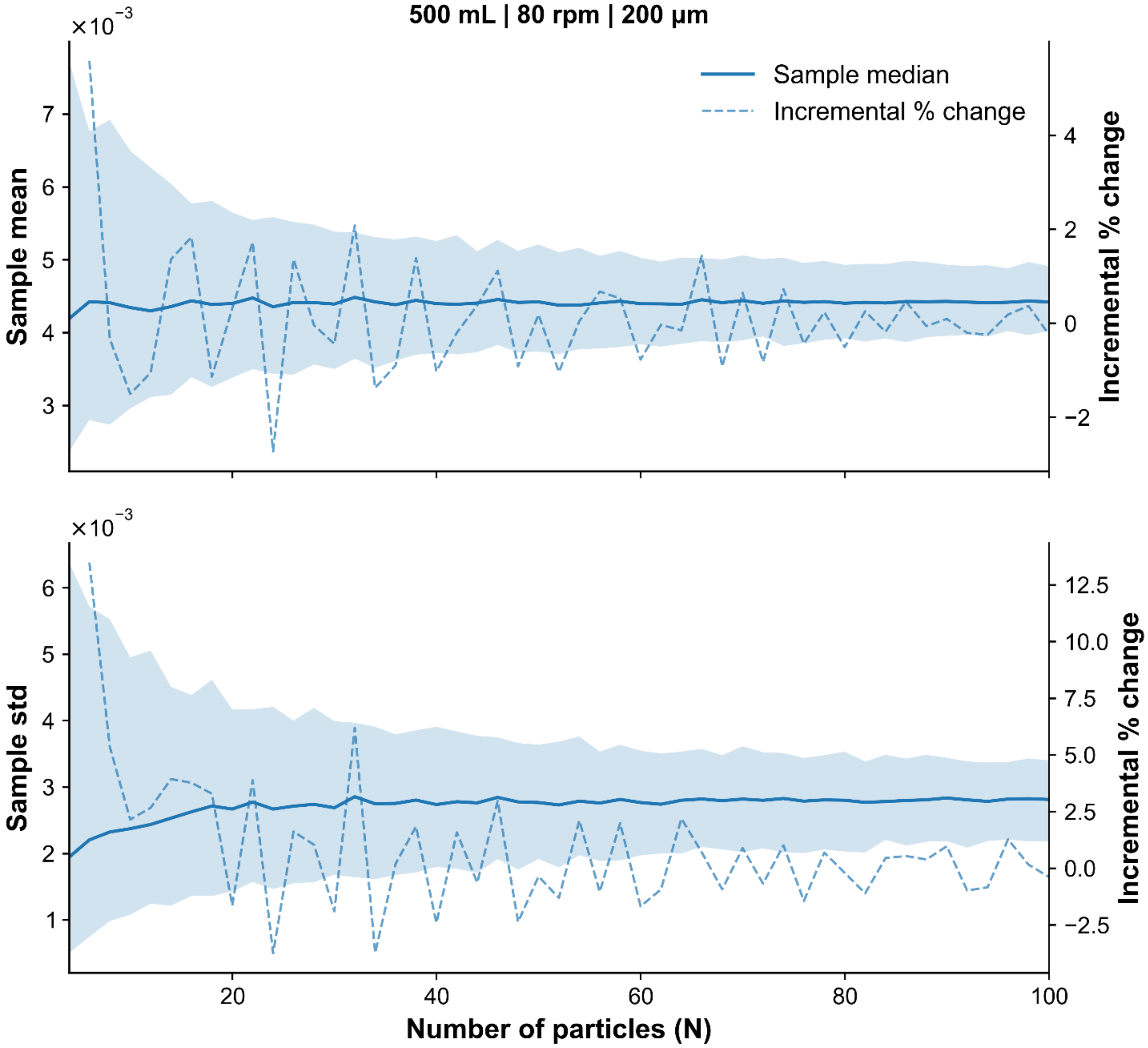


***Fig. S6*** *Particle-number (N) convergence of time-averaged EDR exposure statistics for the 500 mL vertical wheel bioreactor simulated at 80 rpm with 200 μm particles. The solid blue line (median) and shaded band (95% interval) show the across-particle mean (top) and standard deviation (bottom). The 95% interval narrows with increasing N and is tight by N ≈ 30–50 (very tight at N = 100), supporting 100 particles as a conservative sample size. The dashed curve (right axis) shows the incremental % change between successive N values* $[100\,(y_j - y_{j-1})/y_{j-1}]$*, which fluctuates around zero and diminishes as N increases.*

### Particle number independence (Shear, 500 mL VWBR, 80 rpm)

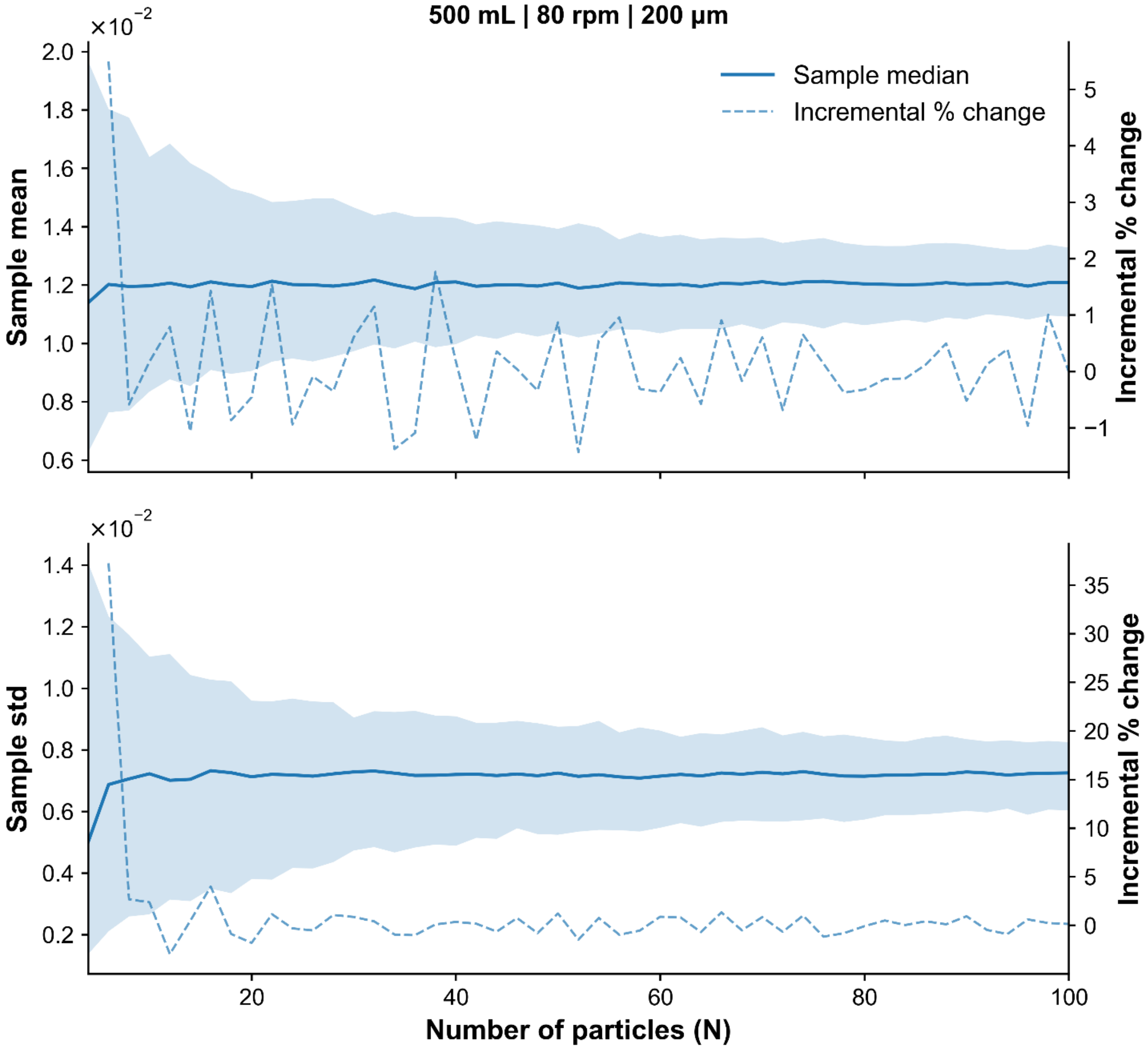


***Fig. S7*** *Particle-number (N) convergence of time-averaged shear-stress exposure statistics for the 500 mL vertical wheel bioreactor simulated at 80 rpm with 200 μm particles. The solid blue line (median) and shaded band (95% interval) show the across-particle mean (top) and standard deviation (bottom). The 95% interval narrows with increasing N and is tight by N ≈ 30–50 (very tight at N = 100), supporting 100 particles as a conservative sample size. The dashed curve (right axis) shows the incremental % change between successive N values* $[100\,(y_j - y_{j-1})/y_{j-1}]$*, which fluctuates around zero and diminishes as N increases.*

#### Tracking time independence (EDR, 100 mL VWBR, 20 rpm)

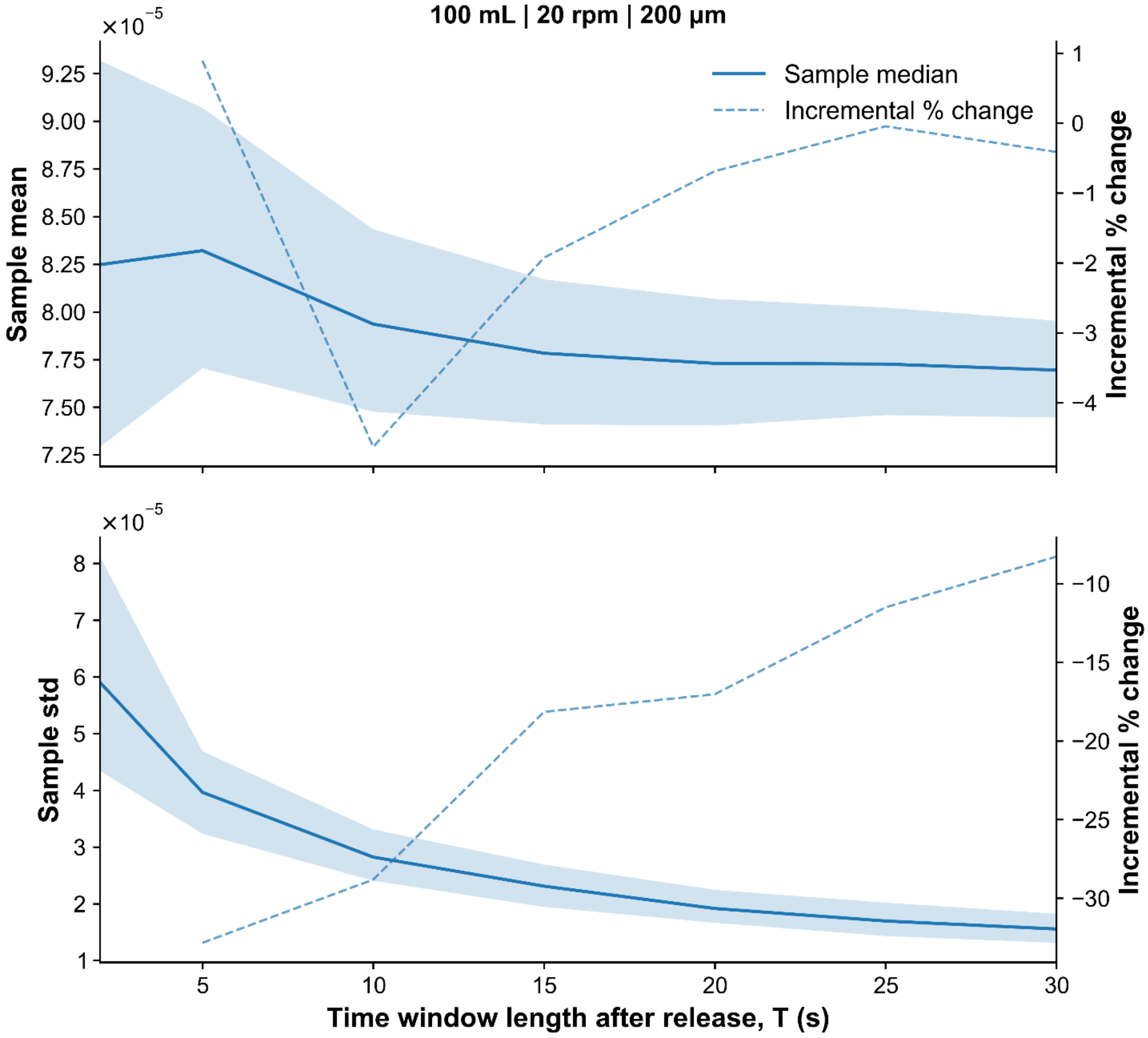


***Fig. S8*** *Time-window (T) convergence of time-averaged EDR exposure statistics for the 100 mL vertical wheel bioreactor simulated at 20 rpm with 200 μm particles. The solid blue line (median) and shaded band (95% interval) show the across-particle mean (top) and standard deviation (bottom) computed over* $[30, 30 + T]$*s after release. Both metrics stabilize by T ≈ 20–30 s, supporting 30 s as an adequate tracking duration. The dashed curve (right axis) shows the incremental % change between successive T values* $[100\,(y_j - y_{j-1})/y_{j-1}]$*, which decreases in magnitude as T increases.*

#### Tracking time independence (Shear, 100 mL VWBR, 20 rpm)

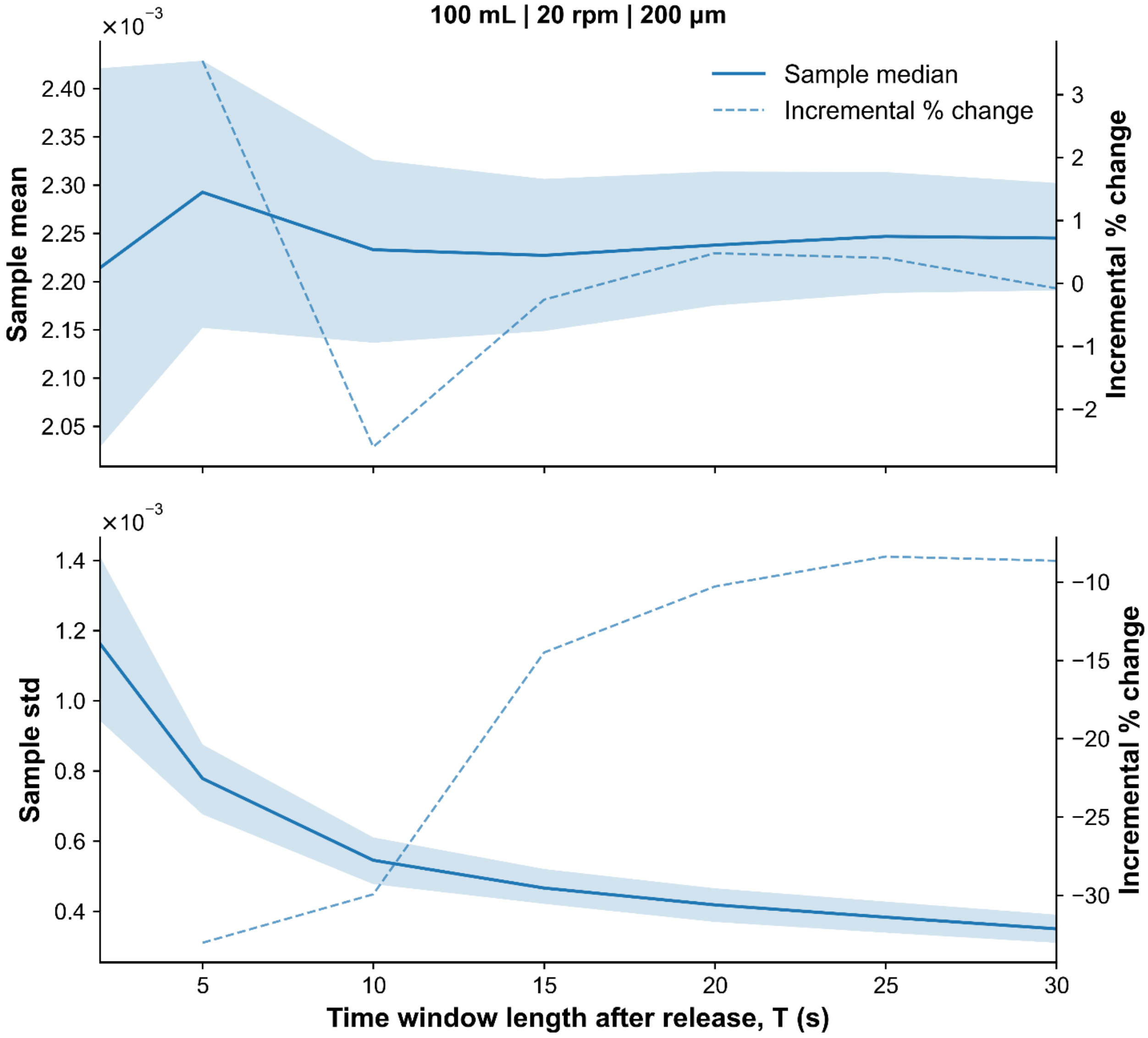


***Fig. S9*** *Time-window (T) convergence of time-averaged shear-stress exposure statistics for the 100 mL vertical wheel bioreactor simulated at 20 rpm with 200 µm particles. The solid blue line (median) and shaded band (95% interval) show the across-particle mean (top) and standard deviation (bottom) computed over* $[30,\ 30+T]$*s after release. Both metrics stabilize by T ≈ 20–30 s, supporting 30 s as an adequate tracking duration. The dashed curve (right axis) shows the incremental % change between successive T values* $[100\ (y_j - y_{j-1})/y_{j-1}]$*, which decreases in magnitude as T increases.*

### Tracking time independence (EDR, 500 mL VWBR, 80 rpm)

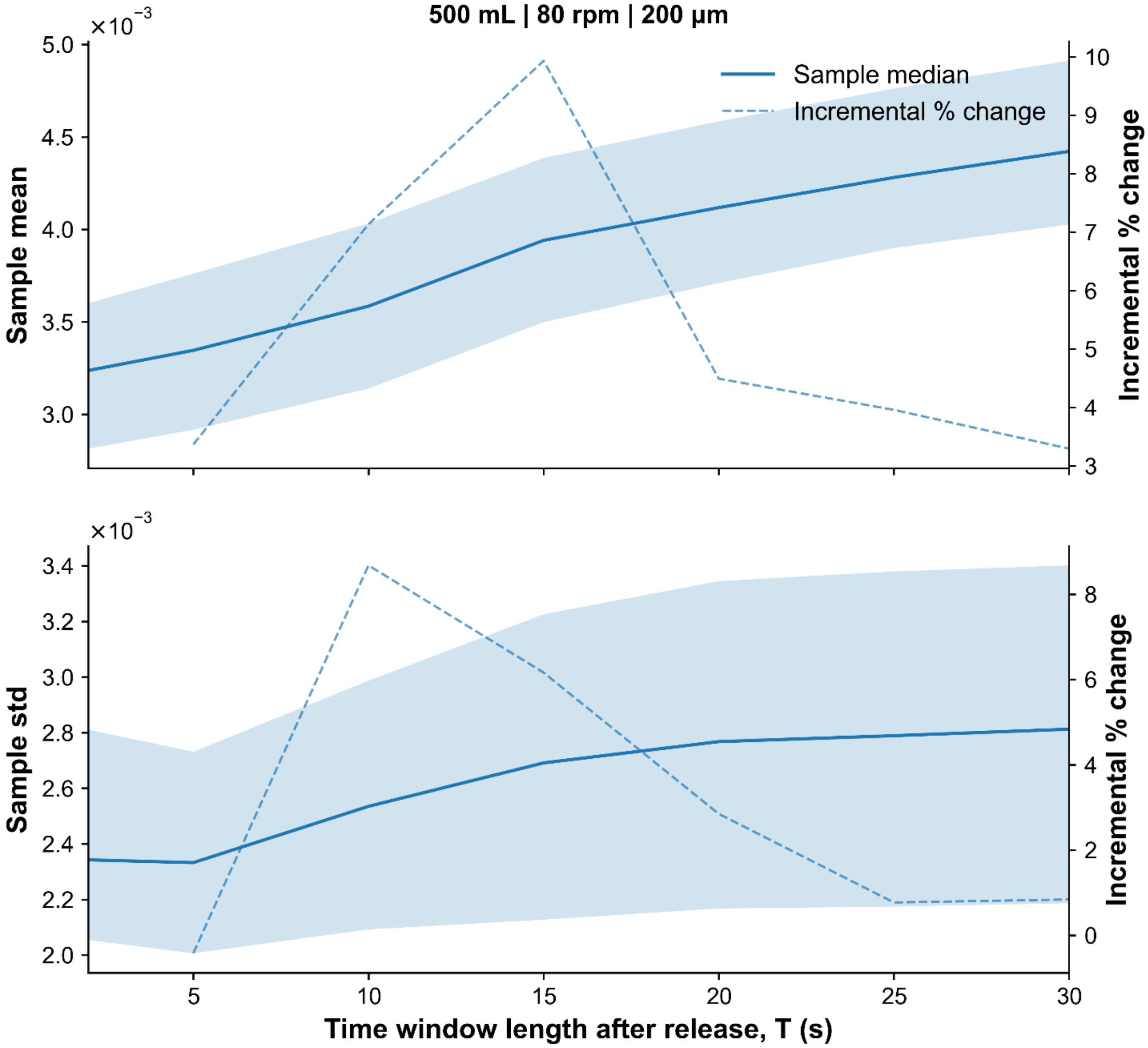


***Fig. S10*** *Time-window (T) convergence of time-averaged EDR exposure statistics for the 500 mL vertical wheel bioreactor simulated at 80 rpm with 200 µm particles. The solid blue line (median) and shaded band (95% interval) show the across-particle mean (top) and standard deviation (bottom) computed over* $[30, 30 + T]$*s after release. The exposure statistics approach a stable range by T ≈ 20–30 s, supporting 30 s as an adequate tracking duration. The dashed curve (right axis) shows the incremental % change between successive T values* $[100\,(y_j - y_{j-1})/y_{j-1}]$, *which decreases in magnitude as T increases.*

#### Tracking time independence (Shear, 500 mL VWBR, 20 rpm)

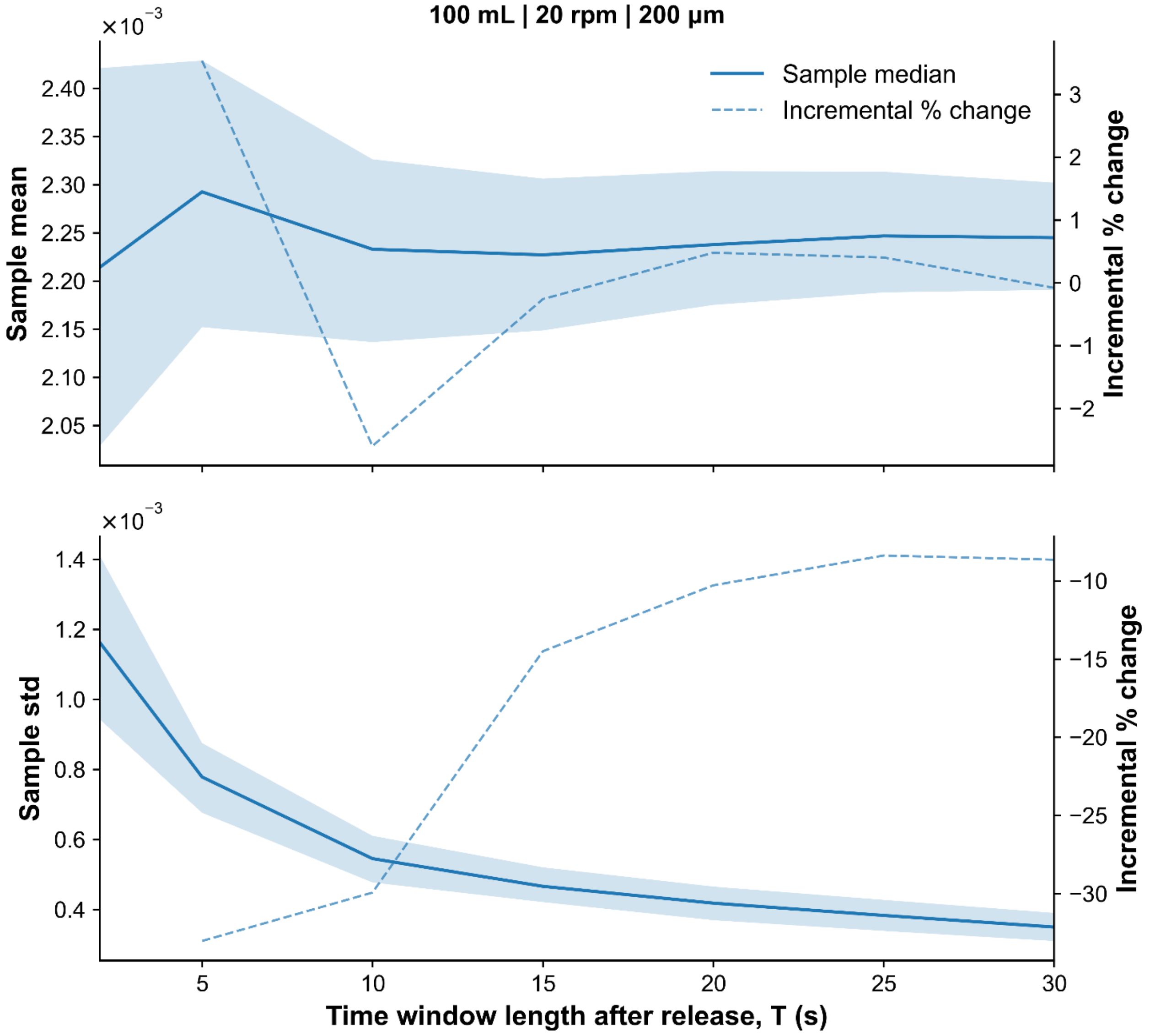


***Fig. S11*** *Time-window (T) convergence of time-averaged shear-stress exposure statistics for the 500 mL vertical wheel bioreactor simulated at 80 rpm with 200 µm particles. The solid blue line (median) and shaded band (95% interval) show the across-particle mean (top) and standard deviation (bottom) computed over* $[30,\, 30+T]$*s after release. The exposure statistics approach a stable range by T ≈ 20–30 s, supporting 30 s as an adequate tracking duration. The dashed curve (right axis) shows the incremental % change between successive T values* $\left[100\,(y_j - y_{j-1})/y_{j-1}\right]$*, which decreases in magnitude as T increases.*

### Average Daily Fold Ratios

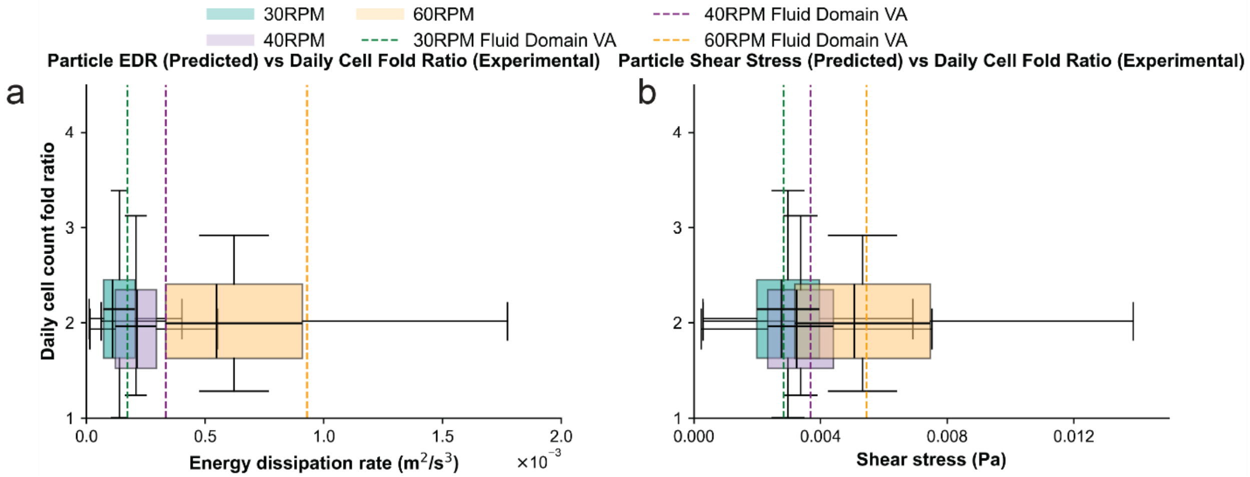


***Fig. S12*** *Average daily fold ratios as a function of EDR (left) and shear stress (right) distributions at various agitation rates in the 100 mL VWBR.*

#### Individual Daily Fold Ratios

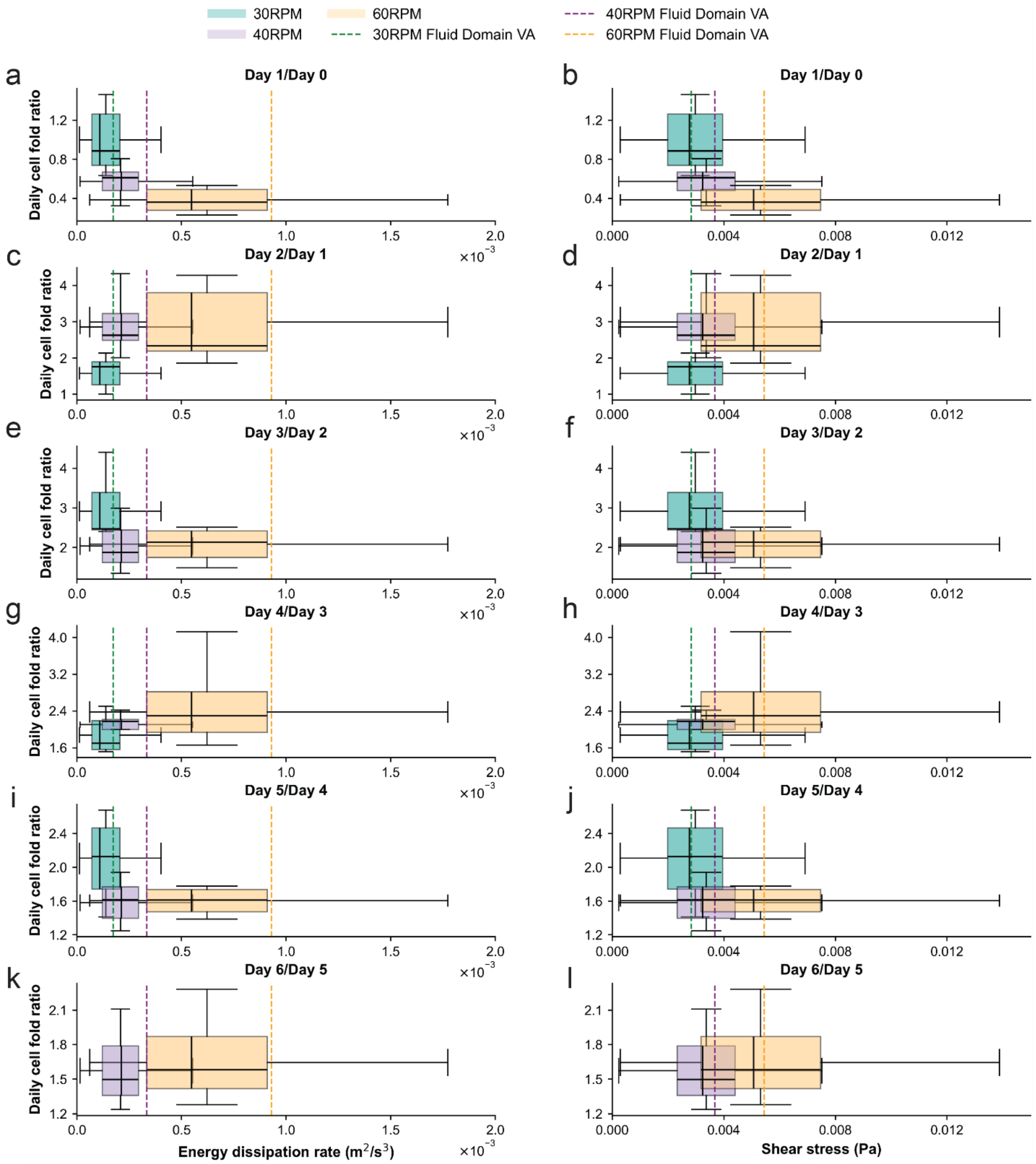


***Fig. S13*** *Daily fold ratios as a function of EDR (left column) and shear stress (right column) distributions at various agitation rates and particle sizes in the 100 mL VWBR. Note that Day 6/ Day 5 did not have cell data available for 30 rpm due to early harvesting of the cells because of cells pooling at the bottom of the bioreactor.*

#### Overall Fold Ratio Final/D2

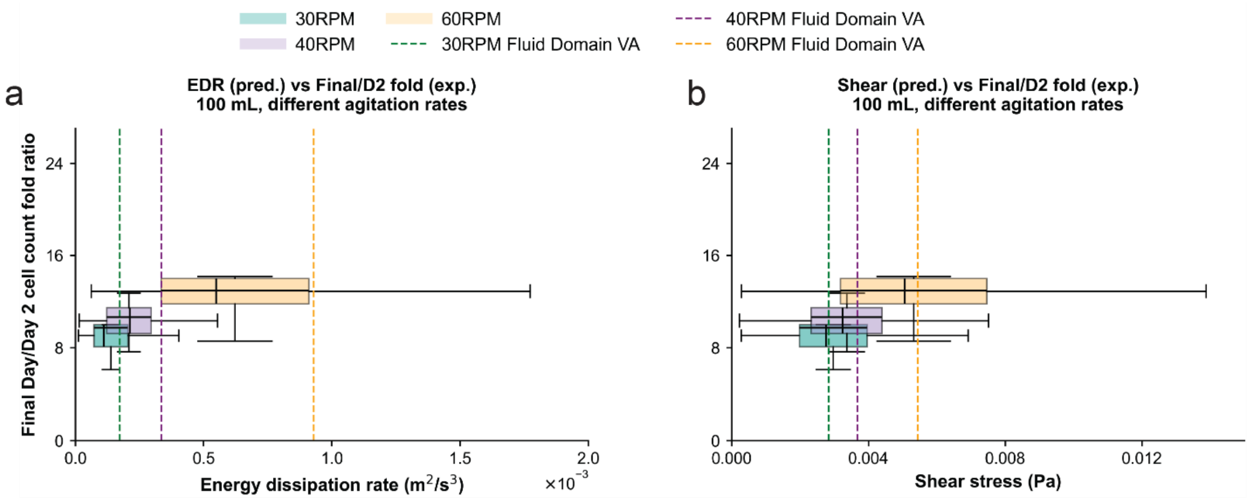


***Fig. S14*** *Final day/Day 2 cell count fold ratios as a function of EDR (left) and shear stress (right) at various agitation rates in the 100 mL VWBR.*

#### Density of the Fluid Region (air, liquid)

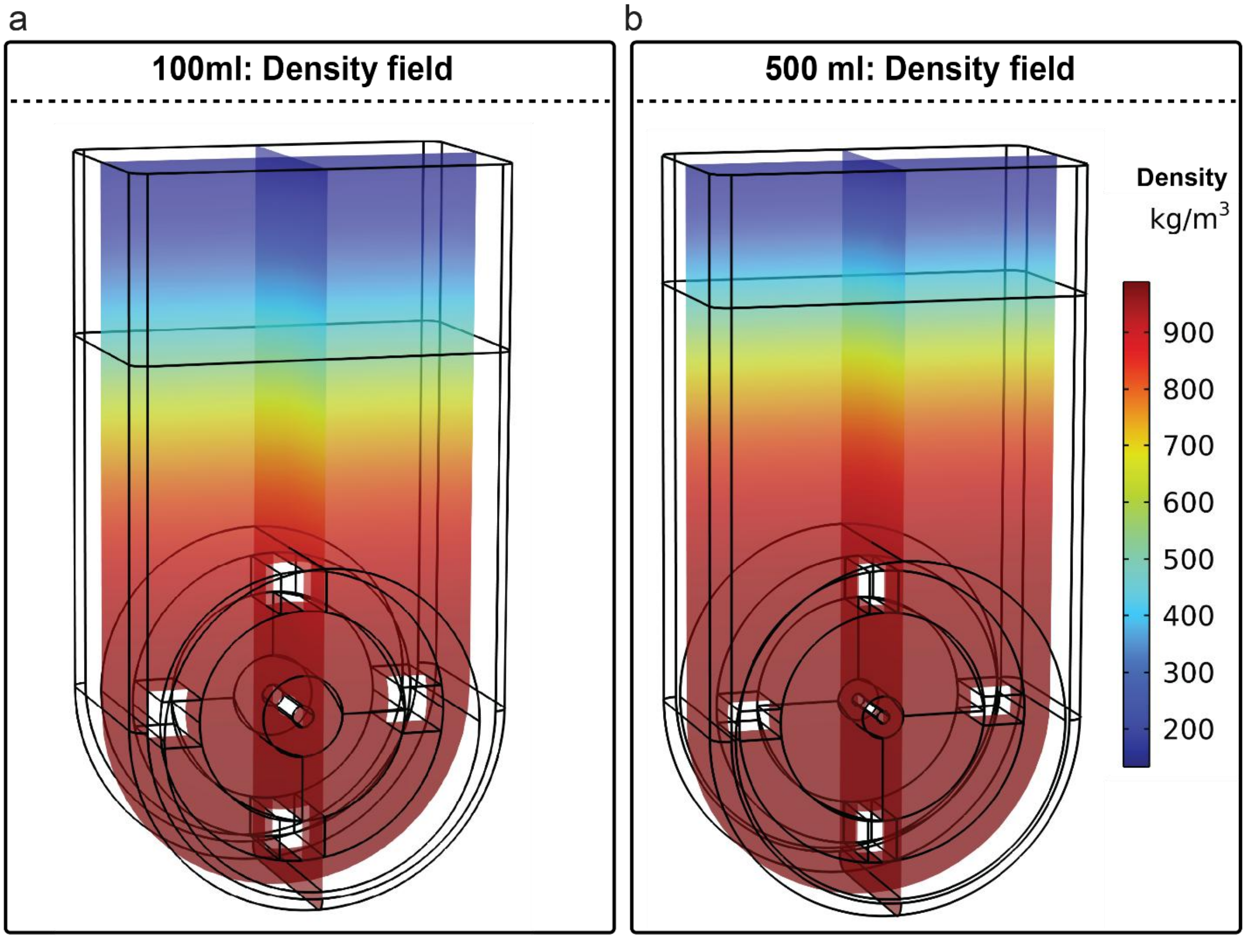


***Fig. S15*** *Two-phase (air–liquid) CFD setup shown via density field. Density (kg/m³) is plotted to visualize the free surface in the vertical wheel bioreactor simulations: low-density air occupies the headspace above the interface, while the high-density culture medium (mTeSR1) fills the liquid domain below. Panels show the two VWBR sizes used in this study: (a) 100 mL and (b) 500 mL. The density contrast confirms the two-phase, free-surface formulation used for the CFD model.*

#### Standard drag correlations

In the particle tracing simulations, the drag force was computed using standard drag correlations available in COMSOL, where the drag coefficient $C_D$ is defined as a piecewise function of the relative Reynolds number $Re_r$.

The relative Reynolds number is

$$Re_r = \frac{\rho \ | \mathbf{u} - \mathbf{v} | \ d_p}{\mu},$$

where $\rho$ and $\mu$ are the fluid density and dynamic viscosity, $d_p$is the particle (aggregate) diameter, $\mathrm{u}$ is the local fluid velocity, and $\mathrm{v}$ is the particle velocity.

Table S1 below shows the drag correlations used, along with the range of particle relative Reynolds number.

***Table S1*** *Standard drag correlations used for particle drag in COMSOL. Piecewise definition of the drag coefficient* $C_D$*as a function of the relative Reynolds number* $Re_r$*, as implemented in COMSOL when the drag law is set to Standard drag correlations. Here* $w = log_{10}(Re_r)$

| Range | Correlation |
|---|---|
| $Re_r \leq 0.01$ | $C_D = \dfrac{24}{Re_r\left(1 + \left(\frac{3}{16}\right) Re_r\right)}$ |
| $0.01 < Re_r \leq 20$ | $C_D = \dfrac{24}{Re_r\left(1 + 0.1315\, Re_r^{0.82 - 0.05\, w}\right)}$ |
| $20 < Re_r \leq 260$ | $C_D = \dfrac{24}{Re_r\left(1 + 0.1935\, Re_r^{0.6305}\right)}$ |
| $260 < Re_r \leq 1500$ | $\log_{10}(C_D) = 1.6435 - 1.1242\, w + 0.1558\, w\hat{}2$ |
| $1500 < Re_r \leq 1.2 \times 10^4$ | $\log_{10}(C_D) = -2.4571 + 2.5558\, w - 0.9295\, w\hat{}2 + 0.1049\, w\hat{}3$ |
| $1.2 \times 10^4 < Re_r \leq 4.4 \times 10^4$ | $\log_{10}(C_D) = -1.9181 + 0.6370\, w - 0.0636\, w\hat{}2$ |
| $4.4 \times 10^4 < Re_r \leq 3.38 \times 10^5$ | $\log_{10}(C_D) = -4.3390 + 1.5809\, w - 0.1546\, w\hat{}2$ |
| $3.38 \times 10^5 < Re_r \leq 4 \times 10^5$ | $C_D = 29.78 - 5.3\, w$ |
| $4 \times 10^5 < Re_r \leq 1 \times 10^6$ | $C_D = 0.1\, w - 0.49$ |
| $1 \times 10^6 < Re_r$ | $C_D = 0.19 - 8 \times \dfrac{10^4}{Re_r}$ |
